# *Toxoplasma gondii* aspartic protease 5 (*Tg*ASP5): Understanding structural details and inhibition mechanism

**DOI:** 10.1101/2023.02.16.528815

**Authors:** Satadru Chakraborty, Anuradha Deshmukh, Pooja Kesari, Prasenjit Bhaumik

**Author notes:** **Corresponding authors**: Pooja Kesari, Department of Biosciences and Bioengineering, Indian Institute of Technology Bombay, Powai, Mumbai-400076, India., Prasenjit Bhaumik, Department of Biosciences and Bioengineering, Indian Institute of Technology Bombay, Powai, Mumbai-400076, India.

## Abstract

*Toxoplasma gondii*, a worldwide prevalent parasite, is responsible for causing toxoplasmosis by infecting almost all warm-blooded animals, including humans. To establish a successful infection, the parasite exports a series of effector proteins which modulates the host immune system; Golgi-resident *T. gondii* aspartyl protease 5 (*Tg*ASP5) plays an essential role in the maturation and export of these effector proteins. This is the first report of the detailed structural investigation of the *Tg*ASP5 mature enzyme. Molecular modeling and all-atom simulation provided in-depth knowledge of the active site architecture of *Tg*ASP5. The analysis of the binding mode of TEXEL substrate highlighted the importance of the active site residues forming the pocket; the Ser505, Ala776 and Tyr689 in the S2 binding pocket provide the specificity towards Arg at the P2 position. Our study also provides insights into the binding mode of the known inhibitor RRL_Statine._ Screening the known aspartic protease inhibitors against *Tg*ASP5 active site and performing 100 ns all-atom molecular dynamic simulations, MM-PBSA binding energy calculations provided the best nine inhibitor protein complexes. Besides that, Principal Component Analysis (PCA) was employed to identify the change in protein dynamics with respect to the substrate and ligand binding. *Tg*ASP5 is essential for the fitness and virulence of the parasite; inhibiting this enzyme can be a possible therapeutic strategy against toxoplasmosis. Our study put forth the inhibitors which can act as initial scaffolds for developing potent mechanistic inhibitors against *Tg*ASP5.

## 1. Introduction

Parasites belonging to phylum apicomplexan are the causative agents of some of the most widespread diseases that are major global health concerns [1]. *Babesia, Theileria, Besnoitia and Eimeria* are associated with pathogenesis in other animals, whereas *Toxoplasma, Plasmodium and Cryptosporidium* cause infection in humans besides infecting other animals and are associated with a huge number of life loss [2–5]. Amongst them, *Toxoplasma gondii* is one of the most widely distributed, causing chronic and acute infections fatal for immunocompromised individuals and newborns. *T. gondii* has a complex life cycle that involves sexual reproduction in feline (definitive host) host, followed by environmental sporozoites which enter into the intermediated hosts (most warm-blooded animals including humans) and disseminates into different host cells as tachyzoites. The rapidly replicating tachyzoites cause acute infection, whereas the slow-growing bradyzoite is responsible for chronic infection resulting in cysts formation in the brain and striated muscle [6,7]. The current therapeutic options are limited to pyrimethamine, trimethoprim, and sulfonamides, primarily work on the de novo folate biosynthesis, which causes significant adverse effects [8–10]; also, increasing reports of drug-resistant strains [11] marks the need for more effective drugs against novel targets. Besides that, *T. gondii* serves as a model for the obligate intracellular apicomplexan parasites [12], inhibitors designed against a novel target for *T. gondii* can also be helpful for designing antiparasitic compounds against other apicomplexans.

Pepsin-like aspartic proteases play vital roles in the life cycle of apicomplexans; ten plasmepsins (Pepsin-like aspartic proteases of *Plasmodium*) express in different life cycle stages of *Plasmodium* which play key roles in necessary life cycle stages- hemoglobin degradation, invasion & egress and host cell remodeling and considered to be important drug targets for *Plasmodium* [13,14]. In *T. gondii* seven pepsin-like aspartic proteases (*Tg*ASPs) have been identified, which express in acute infection causing tachyzoite stage and cysts forming bradyzoite stage [15]. Amongst them *Tg*ASP3, and *Tg*ASP5 have been characterized recently, which play necessary roles in invasion & egress of *T. gondii* and host cell remodeling respectively [16,17] and essential for parasite survival and virulence.

*T. gondii* modulates the host immune response to remain undetected and survive inside the host cell, even for a lifetime. This modulation is done by a series of dense granular proteins transported towards the host cell, crossing the parasitophorous vacuole membrane (PVM). After the invasion of tachyzoite and bradyzoite inside host cells, the parasite stays in the parasitophorous vacuole (PV), and PVM separates the parasite from the host cell cytoplasm [18]. The *T. gondii Tg*ASP5 serves as a critical player in the protein export pathway, responsible for the priming of dense granular proteins (GRAs) before exporting through PVM into the host cell. GRA proteins possess a pentameric motif called TEXEL or *Toxoplasma* EXport ELement [17,19]. *Tg*ASP5 shares a sequence identity of 31.61% with plasmepsin V (PMV) of *Plasmodium vivax*, which is found in the endoplasmic reticulum of *Plasmodium* and is responsible for cleaving the PEXEL (*Plasmodium* EXport ELement) motif at RXL↓XE/Q/D (X denotes any amino acid) [20] site in the proteins before exporting from PVM. However, *Tg*ASP5 is a Golgi resident protease that cleaves substrates at RRL↓XX. Unlike the PEXEL motif, the substrate of *Tg*ASP5 requires Arg at P2 position which is necessary for substrate binding and catalysis. Though PEXEL is present only in the N-terminal of the substrates, TEXEL can be at any position. The enzymes also get inhibited by the substrate mimetic inhibitor RRL_Statine,_ which contains Arg at P3 and P2 positions, while RVL_Statine_, a PEXEL mimetic, cannot inhibit the enzyme [17]. These PEXEL-like motifs were also found in secreted proteins of *Babesia bovis* and *Cryptosporidium parvum* [21]. Plant pathogen oomycetes also contain a similar RXLR motif required for targeting effector proteins towards the host cell [22].

Besides processing proteins with TEXEL motif, *Tg*ASP5 is also involved in exporting some GRAs that do not possess the TEXEL motif [17]. It is associated with processing proteins (MYR1, MYR4, GRA44, GRA45), which are part of the transportation machinery of GRAs [23]. The deletion of *Tg*ASP5 decreases the virulence and fitness of the parasite; severe attenuation of this *Tg*ASP5 deficient *Toxoplasma* was observed in the murine model. In absence of *Tg*ASP5 the GRAs cannot cross the PVM, membranous nanotubular network (MNN) formation, host cell mitochondrial recruitment got impaired, global changes of gene expression happen and the parasite becomes more susceptible to the host’s immune system [17,19].

The important role of *Tg*ASP5 in protein export, involved in host cell remodeling, made it a potential drug target for toxoplasmosis. For designing a potent and selective inhibitor against the enzyme, a detailed understanding of the substrate binding pocket is required. In the absence of the experimental structure, we have developed a structural model of *Tg*ASP5, elucidating the important structural features and identifying key active site residues required for interactions with substrate. The molecular basis of binding of the TEXEL substrate is being investigated in detail with the help of docking and molecular dynamics simulation. The binding mode of only reported *Tg*ASP5 inhibitor RRL_Statine,_ which is a TEXEL mimetic, is being predicted. The known mechanistic inhibitors of aspartic proteases were screened, and the binding efficiency was calculated based on molecular dynamics (MD) simulation and Molecular Mechanics Poisson-Boltzmann Surface Area (MM-PBSA) binding energy calculations method to rank the molecules. The best hits from the screened library can act as the initial scaffold for designing highly potent and selective inhibitors against toxoplasmosis. This study provides the molecular basis for designing inhibitors against pepsin-like aspartic proteases of pathogenic apicomplexan parasites.

## 2. Materials and methods

### 2.1 Sequence analysis of *Tg* ASP5

A total of 8 different *Tg*ASP5 sequences were obtained from ToxoDB database (https://toxodb.org/) with ToxoDB entry TGRH88_029500 (*T. gondii RH88*), TGGT1_242720 *(T. gondii GT1)*, TGRUB_242720 (*T. gondii RUB*), TGP89_242720 (*T. gondii p89*), TGME49_242720 (*T. gondii ME49*), TGVAND_242720 (*T. gondii VAND*), TGBR9_242720 (*T. gondii TgCATBr9*), TGARI_242720 (*T. gondii ARI*). The mature segments of the proteins were identified by comparing the sequences of *Tg*ASP5s with human pepsin (UniProt Entry: P0DJD7) and *Plasmodium falciparum* plasmepsin II (*Pf*PMII) (UniProt Entry: Q8I6V3) mature enzyme, which have a characteristic DTGS/D(T/S)GT motif in the N and C terminal, respectively. The mature segment of pepsin-like aspartic proteases usually starts at the 31 amino acids upstream position of the first catalytic aspartate, which is referred to as Asp32 (numbering as per pepsin) [24,25]. Comparing *Tg*ASP5 with these two enzymes suggests that the first catalytic aspartate is at 431th position of the full-length *Tg*ASP5_RH88 sequence. Therefore, the mature portion of *Tg*ASP5 was considered from the residue region Arg400-Phe809 for molecular modeling studies. *Tg*ASP5 like protein sequences from other organisms-*Plasmodium falciparum* PMV (Q8I6Z5), *P. vivax* PMV (Q6PRR9), *P. knowlesi* PMV (A0A5K1US84), *P. ovale* PMV (A0A1A9AK19), *P. malarie* PMV (A0A1A8WW61), *Thioleria orientalis* Aspartic protease 5 (ASP5) (A0A2T7IQU4), *Besnoitia besnoiti* ASP5 (A0A2A9ML77), *Cyclospora cyatensis* ASP5 (A0A1D3D9S0), were also obtained from UniProt database and were aligned with *Tg*ASP5 using Clustal Omega [26,27].

### 2.2 Homology Modeling of mature part of *TgASP5*

Homology modeling was done using SWISS-MODEL [28] based on the template identified by PSI-BLAST as well as SWISS-MODEL template search. The template used for modeling the mature part of *Tg*ASP5 was the crystal structure (PDB ID: 4ZL4, resolution 2.37Å) of plasmepsin V of *P. vivax* with a sequence identity of 36.4%. This initial model was re-iterated for multiple cycles to generate a final model of *Tg*ASP5. The optimization of the model was required because of the varying lengths of the insertion region between the *Pv*PMV and *Tg*ASP5. Phobius [29] (Fig. S1A) and TOPCONS [30] predictions (Fig. S1B) were used to identify transmembrane domains of *Tg*ASP5. The quality of the model was analyzed by Ramachandran plot generated PDBsum [31] and also Verify-3D [32] and ERRAT [33] map of the SAVES server. Detailed structural analysis was done by comparing the newly generated model with other pepsin-like aspartic protease structures by using PyMol [34] and COOT [35].

### 2.3 Preparation of the Substrate

To study the binding mode of the substrate in the *Tg*ASP5 active site, the TEXEL motif sequence ^P3^RRLAE^P2^**’**of the substrate GRA21 of *T. gondii* [17] was used. The structure of RRLAE was prepared by using WEHI-842 (Ligand ID: 4PK) (PDB ID: 4ZL4) as a template. 4PK is a peptidomimetic inhibitor of plasmepsin V (PMV), which mimics the PEXEL sequence RxLxE/Q/D [36]. The designed substrate was docked into *Tg*ASP5 active site to generate a substrate-bound conformation of *Tg*ASP5.

### 2.4 Docking of Substrate and Inhibitors

The known inhibitors of pepsin-like aspartic proteases and HIV-1 proteases were retrieved selected from Protein Data Bank (http://www.rcsb.org). The coordinates of these 80 ligands were manually checked before docking. The coordinates of *Tg*ASP5 and ligands were then converted to pdbqt format using AutoDockTools-1.5.6 [37]. Polar hydrogen and Kollman charges were added with the protein and atoms were assigned as AD4 types before converting to pdbqt files. Gasteiger charges were added with the ligands and detection of root and selection of rotatable and non-rotatable bond was done before converting it into pdbqt format. The grid was selected based on 4PK after superposing the 4PK bound *Pv*PMV with *Tg*ASP5. The grid spacing was kept at the default 0.375, and the final grid dimension was 46×48×46 with grid center −3.040, 99.280, and 42.328. This dimension was finalized after multiple docking runs with 4PK. The grid parameter file was generated and Autogrid4 was run to generate the glg file. The docking was done using Autodock4 [37], and the Genetic Algorithm was used to generate 50 poses. The selection of the relevant poses was made from these 50 conformations. Further, the optimized grid was used for docking other ligands as well as substrate.

### 2.5 Molecular Dynamics (MD) Simulation of the Apo-structure, Substrate and Ligand Bound Structure of *Tg*ASP5

Molecular dynamics (MD) simulation of all *Tg*ASP5 structures was performed using AMBER99SB [38] force field by GROMACS 2019.1 [39]. ‘xleap’ module of AmberTools 2018 [40] was used for generating the topology for the apo structure as well as substrate and ligand bound complexes. The disulfide bonds were appropriately defined before the generation of the topology file. Explicit water model TIP3P or Transferable Intermolecular Potential 3P, a three-point rigid water model, was used for solvation. The cubic solvation box had a distance of 20Å from the protein to the boundary of the box and the counter ions were added to neutralize the systems. The ParmED (https://parmed.github.io/ParmEd) package was imported in Python3.7 and used to convert the topology generated by AmberTools 2018 to GROMACS compatible file formats. The energy minimization was done using the steepest descent algorithm for 50000 steps with a cutoff of 1000 kJ mol^−1^nm^−1^. The neutralized solvent system containing the apo structure was heated to 300 K for 100ps with 2fs time step during constant volume equilibration process. It was followed by equilibration process using an isothermal-isobaric ensemble with Parrinello-Rahman barostat for 100ps with 2fs time step to maintain 1 bar pressure constantly. The production MD simulation was performed for 100 ns for the apo structure with a time step of 2fs. The temperature was maintained at 300 K using v-rescale temperature coupling method with time constant of 0.1 ps. The pressure was maintained at 1 bar by Parrinello-Rahman pressure coupling method with a time constant of 2ps. Lincs algorithm was used to constrain the covalent bonds involving hydrogen. The short-range van der Waals Interactions were calculated for atom pairs using the Verlet scheme with a cut-off of 1nm and for short-range electrostatic interaction it is also 1nm. The particle-mesh Ewald method was used for long-range electrostatic interaction with grid spacing of 1.6Å grid spacing and 4^th^ order cubic interpolation. The trajectories generated from the final MD run were analyzed by GROMACS 2019.1 and VMD (Visual Molecular Dynamics) [41]. The figures were generated by PyMol and Discovery studio visualizer (https://www.3ds.com/products-services/biovia/). The simulation of the substrate-protein complex was performed similarly with the apo structure.

For the simulation of the ligand bound structure the parameters for the ligand file was prepared first. The addition of hydrogen with ligand was done using Chimera 1.14 [42] or Avogadro [43]. After the valency of the atom and addition of hydrogen with ligands were properly defined, AM1-BCC method of the *antechamber* program of the Amber Tools 2018 was run. Van der Walls and bonded parameters for ligands were obtained from General Amber Force Field (GAFF). The complete Amber coordinate files for the ligands were generated using *parmchk2* program. The topology of the ligand bound *Tg*ASP5 complexes were generated ‘xleap’. Similar protocol was followed for performing MD for other complexes as well. The other methods were done similarly to the apo structure of the *Tg*ASP5. To check the structural stability after the final production run, root mean square deviation (RMSD) analysis was done based on backbone after backbone based least square fitting using gmx_RMS_ tool. For analyzing the flexibility of the residue root mean square fluctuation (RMSF) analysis based on C-alpha (CA) atoms, was done on apo, substrate bound, ligand bound structures using gmx_RMSF_ tool of GROMACS. While the radius of gyration was checked for measuring the compactness of the structures using gmx_gyrate_.

### 2.6 MM_PBSA Binding Energy Calculations

The binding energy of ligand bound complexes of *Tg*ASP5 was calculated using g_mmpbsa tool [44]. The different energy components-E_MM_, G_nonpolar_, G_polar_ were calculated for 500 snapshots for the most stable region in the last 50ns simulation. E_MM_ includes both electrostatic (E_elec_) and van der Walls (E_vdW)_ components. The final binding energy is the summation of E_MM_, G_nonpolar_, G_polar._ The 5ns stretches of trajectories containing the 500 snapshots were selected for the region whether RMSD is stable for 500 snapshots. For calculation of G_nonpolar_ solvent accessible surface area (SASA) only model was used.

### 2.7 Principal Component Analysis

Principal Component Analysis (PCA) was done by using the CA atom of the trajectories and by building a covariance matrix using gmx_covar_ tool of Gromacs. To view the movement along PC1 and PC2 gmx_anaeig_ was used. Modevectors.py script was used in pymol to see the detailed vector movement of the residues.

## 3. Results & Discussion

### 3.1 Overall Structural fold of Apo-*Tg*ASP5

The structure of *Tg*ASP5 described in this study was modeled using the sequence of *T. gondii* RH88 (ToxoDB ID: TGRH88_029500); which is a highly virulent strain of *T. gondii* [45,46]. *Tg*ASP5 is a membrane-anchored protease like plasmepsin V [36, 47]; and three transmembrane regions (231-251, 276-296 and 864-884) of the polypeptide were identified by Phobius and TOPCONS (Fig. S1A). The quality of the model of mature *Tg*ASP5 (Arg400-Phe809) was checked by the Ramachandran plot in which 86.5% residues were found to be in the most preferred region, 10.9% and 2.1% residues were found in the additionally allowed and generously allowed regions, respectively; and only 0.6% residues were in the disallowed regions (Fig. S2A). The verify 3D plot suggested 86.59% of the residue to have a score of >= 0.2 in the 3D/1D profile which should be >80% for a proper model. The quality factor of the *Tg*ASP5 model was found to be 86.15%, suggested by ERRAT, which is also in the permissible range. The stability of the model of *Tg*ASP5 was assessed by molecular dynamics (MD) simulation of the structure for 100 ns. The RMSD plot (Fig. S2C) suggests that the structure of *Tg*ASP5 remains stable throughout the last 70 ns of the simulation run. The compactness of the structure is maintained throughout the simulation process, as indicated in the radius of gyration plot (Fig. S2B). The final 100ns simulated structure was taken for further structural analysis.

The structural model of *Tg*ASP5 monomer consisting of 410 amino acids (Arg400-Phe809) can be divided into two domains (Fig. 1A). The N-terminal domain is composed of Arg400-Ser648 and Lys787-Phe809, and the C-terminal domain contains Thr654-Leu781. The substrate binding site is present in-between these two domains. The active site is composed of the characteristic DTGS/DSGT catalytic motifs present at 431-434 in N-terminal and 682-685 in the C-terminal (Fig. 1B) domains. The residues Met499-Val518 are part of the flap region (beta-hairpin) that is placed perpendicular to the active site pocket and is involved in substrate binding. Generally, two disulfide bonds are present in vacuolar plasmepsins [25], and three are present in pepsin and plasmepsin X mature structures [14,48]. However, in the structure of *Pv*PMV and *Tg*ASP5, a total of seven disulfide bonds are present in the mature segment. The Cys441-Cys538, Cys444-Cys447, Cys474-Cys498, Cys468-Cys478 and Cys587-Cys805 are present in the N-terminal domain whereas, Cys706-Cys712 and Cys721-Cys766 reside in the C-terminal domain of the *Tg*ASP5 structure (Fig. 1C). The positions of these 14 cysteine residues, which are engaged in disulfide bonds in *Tg*ASP5, are also conserved in other ASP5 enzymes (Fig. S3).

**Figure 1.**
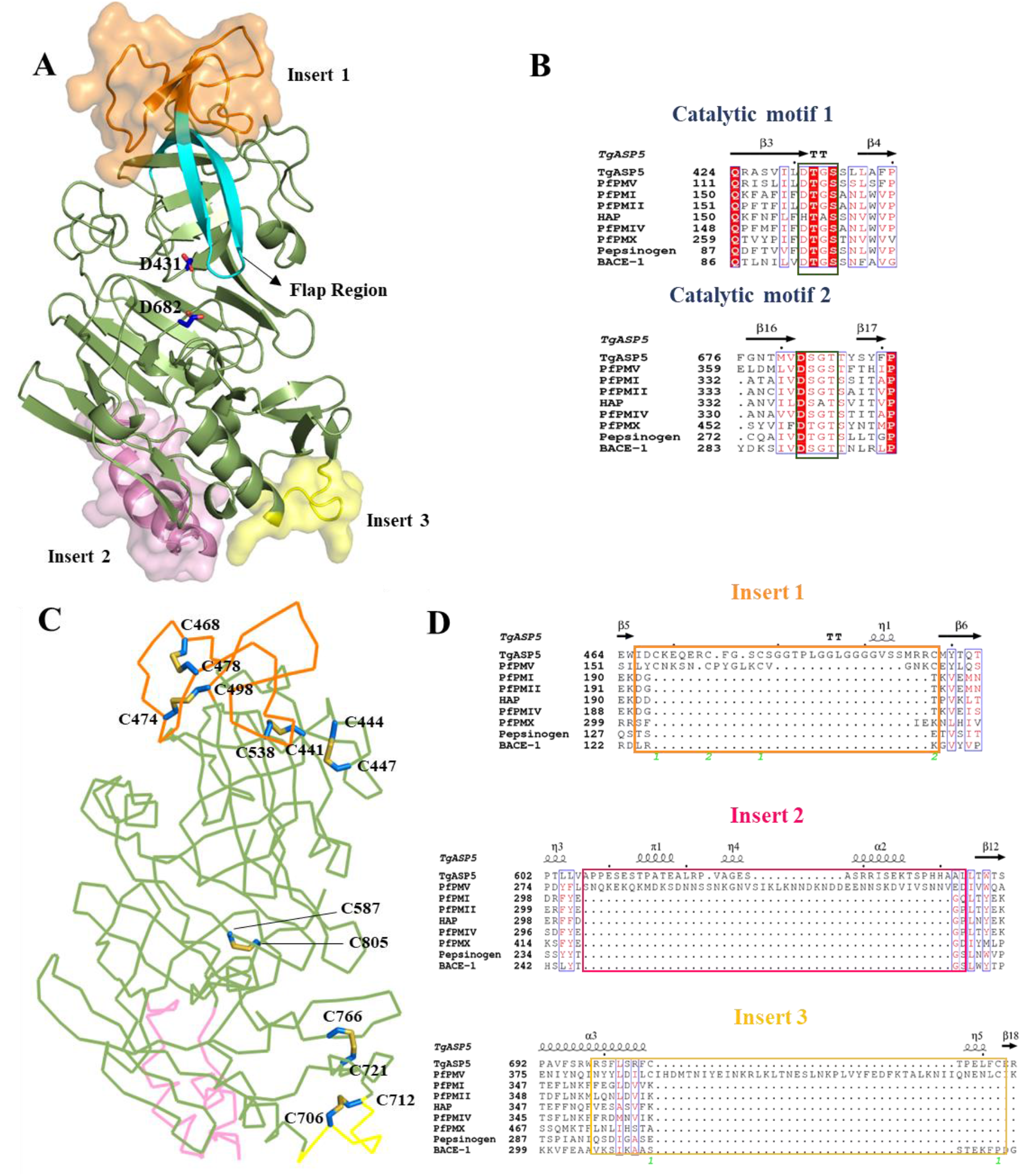
Overall structural feature of ASP5 of *Toxoplasma gondii* (*Tg*ASP5). **(A)** The structure of mature *Tg*ASP5 is represented in the cartoon form. The classical pepsin-like aspartic protease fold is shown in smudge green and flap in cyan, Insert-1 (NAP-1), Insert-2 and Insert-3 are shown in semi-transparent surfaces in orange, pink and yellow color respectively. The catalytic aspartates are represented as royal blue sticks. **(B)** Presence of the two conservative catalytic motifs of pepsin-like aspartic proteases. The identical residues of the alignment are shown in white color in red background and similar residues are represented in red color generated by ESPript (https://espript.ibcp.fr/ESPript/ESPript/). The secondary structural features are of *Tg*ASP5 mature enzyme. **(C)** The presence of seven disulfide bonds is shown in pearl blue sticks, the enzyme is in ribbon representation. The color coding is the same as panel (A). **(D)** Alignment of the insert regions of *Tg*ASP5 with known aspartic proteases.

### 3.2 Comparison with other pepsin-like aspartic proteases

The structural superposition (SSM superpose) of the *Tg*ASP5 model with the crystal structure of *Pv*PMV (PDB ID: 4ZL4) showed an overall RMSD of 2.47 Å (Fig. S1B). Whereas, the structural superposition with other pepsin-like aspartic proteases produced RMSD values of 2.52 Å, 2.64 Å, 2.86 Å, 2.9 Å, 2.9 Å, 3.12 Å with Uropepsin (PDB ID: 1FLH), human Beta-secretase 1 (PDB ID: 3IXJ), Human pepsin (PDB ID:1PSO), mouse submaxillary renin (PDB ID: 1SMR), *P. falciparum* plasmepsin II (PDB ID: 1LF3) (Fig. S1C), *P. falciparum* plasmepsin I (PDB ID: 3QRV), respectively. These values indicate that *Tg*ASP5 has a pepsin-like aspartic protease structural fold, which is very similar to the fold observed among this A1 family of aspartic proteases.

Three unique insertions are found in the sequence of *Tg*ASP5 and plasmepsin V, which are not observed in pepsin and other vacuolar plasmepsins (Fig. 1D). In *Tg*ASP5, Insert-1 contains 33 amino acids (Ile466-Cys498), Insert-2 has 35 residues (Ala607-Leu643) and Insert-3 has 8 amino acids (Arg704-Cys712). These insert regions of *Tg*ASP5 and PMV from *Plasmodium* are of variable lengths. The Insert-1 of *Pf*PMV (Leu153-Cys171) has 19 residues [49] instead of 33 of *Tg*ASP5. The extra stretch of the residues in Insert-1 of *Tg*ASP5 is rich in glycine residues (Fig.1D, S4). This Insert-1 is referred to as Nepenthesin I (Nep I) like sequence, a signature of the A1B family of proteases. Two disulfide bonds (Cys468-Cys478) and (Cys474-Cys498) (Fig. 1C) are present in the Insert-1 and these bonds are known to provide stability over a wide range of temperature and pH condition in the nepenthesin-like enzymes [50,51]. The Insert-I disulfide bonds also hold the tip of the β-hairpin structure of the flap in place. The Insert-2 (Leu278-Val326) of *Pv*PMV forms a helix-turn-helix fold [49], while in *Tg*ASP5 (Ala607-Leu643) it is a small helix. The Insert-3 (Ile387-Cys434) is the key structural feature of *Pv*PMV, present in all *Plasmodium* species, and contains a helix-turn-helix fold [36]; however, this region is much smaller in *Tg*ASP5 (Arg704-Cys712) and other ASP5 enzymes (Fig. S3), and contain a loop region (Fig. 1A). The α3 helix present before Insert-3 of *Tg*ASP5 and PMV is longer, compared to plasmepsin II (Fig. S1C, S3). The Insert-1 and Insert-2 show huge fluctuation compared to the overall structure (Fig. S2D).

### 3.3 Active site architecture of apo-*Tg*ASP5

The active site of the *Tg*ASP5 cleft (Fig. 2A) is formed by the catalytic motifs (^431^DTGS^434^ and ^682^DSGT^685^) on two sides and the flap covering it from the top. The conformational flexibility of the residues lying the active site cleft of *Tg*ASP5 was monitored throughout the MD simulation run to assess the compactness of the region. The distance between the CG atom of the two key catalytic aspartates Asp431 and Asp682 was maintained at 6Å-8Å (Fig. 2B), enabling water-mediated interactions. The density of water molecules around both the catalytic aspartates was also monitored and the data shows the presence of water molecule within 2.5Å of the C-terminal Asp (Fig. 2C). The water molecule near the C-terminal catalytic aspartate produces the hydroxyl anion which acts as a nucleophile to initiate the catalytic reaction [52]. The Asp431 forms a hydrogen bond with Ser434, and this interaction helps the catalytic Asp to remain in protonated form. The hydrogen bond between the side chain of Thr685 and Asp682 is also a key interaction and necessary for the stability of the active site. Both these interactions with catalytic Asp are maintained for complete 100 ns simulation (Fig. 2D, 2E). In classical pepsin-like aspartic protease, the Tyr of the flap forms a hydrogen bond interaction with Trp39 and water mediated interaction with Ser35 in the presence of substrate or ligand, for an extended hydrogen bonding network considered to be necessary for catalysis [52]. In *Tg*ASP5, the flap Tyr is at 504^th^ position while Trp is replaced by Ala438. The Tyr of the flap is conserved in the ASP5-like proteases belonging to the A1B subfamily but Trp39 is replaced by Ala, Ser, Asn (Fig. S3), which indicates these enzymes might have a modified catalytic mechanism compared to pepsin and pepsin-like enzymes.

**Figure 2.**
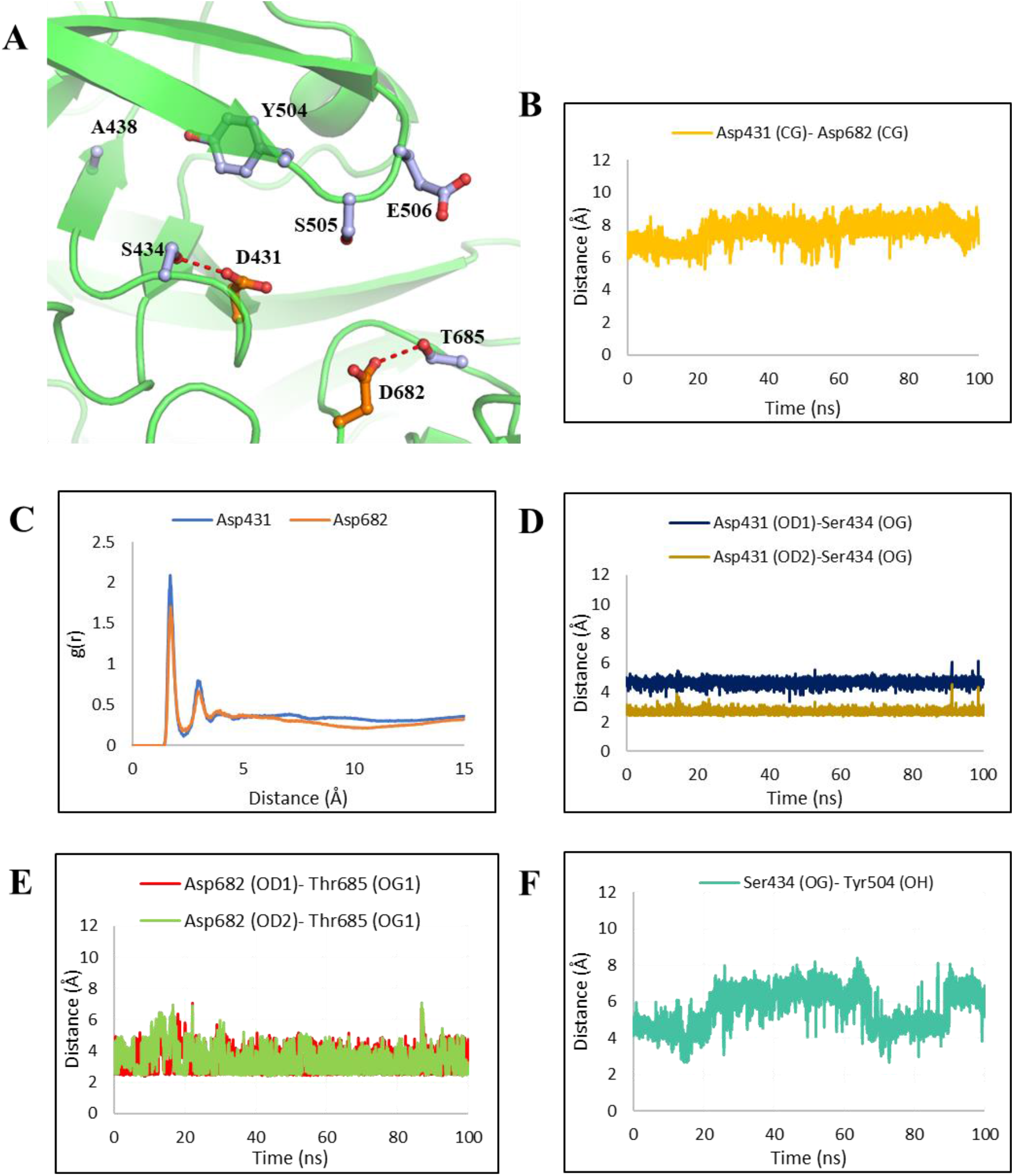
Active site architecture of *Tg*ASP5 apo structure. **(A)**The organization of key active site residues of *Tg*ASP5 involved in catalysis. The two catalytic residues are shown in orange (carbon) ball and sticks while other important residues are represented in light blue (carbon) ball and sticks. The hydrogen bonding networks are marked with red dashes. **(B)** The distance between CG atoms of the Asp431 and Asp682 during the simulation run, capable of containing catalytic water molecule. **(C)** The radial distribution function g(r) of the water molecule hydrogen atoms around the OD1 and OD2 of the Asp431 & Asp682. **(D), (E)** Presence of stable hydrogen bonding interaction between OG atom of active site Ser434 and OG1 of Thr685 with catalytic OD1 & OD2 atoms of Asp431 and Asp682, respectively. **(F)** Change in distance between OG atom of active site Ser434 with OH atom of flap Tyr504 during the simulation process, depicting the flap movement.

The flap is fluctuating during the simulation process as observed from the change in the distance between OG atoms of Ser434 and the OH atom of Tyr504 of the flap (Fig. 2F) and CA atom of Ser505 of the flap tip and CA atom of Asp431 (Fig. S5A). In *Pv*PMV crystal structure, the CA atom of Cys140 of the flap tip and the CΑ atom of catalytic Asp80 are present at 12.4Å apart. In *Tg*ASP5, at initial 4 ns the CA of Ser505 of the flap and CA atom of catalytic Asp431 were at 8.8Å distance (Fig. S5B), the close conformation of the flap which will probably not allow binding of the substrate to the active site. After 26 ns the flap opens to 13.4Å (Fig. S5C), making this pocket accessible to incoming substrate. However, after 52 ns the distance becomes 10.5Å (Fig. S5D). Besides the flap, the flexible loop region (Asn772-Ala776) which is present exactly opposite to the flap above the C-terminal catalytic motif shows dynamic behavior, also associated with opening and closing of the active site pocket (Fig. S5B, S5C, S5D).

### 3.4 Mode of binding of the substrate in the active site of *Tg*ASP5

TEXEL sequence motif is observed at the cleavage site of *Tg*ASP5 substrates [17]. To study the interactions of the TEXEL motif containing substrates with *Tg*ASP5, a peptide substrate was docked into the active site and simulated. The substrate sequence used for the study was ^P3^RRL/AE^P2́^ which is the cleavage site sequence of dense granular protein 16 (GRA16, 63-67 amino acid position) and GRA21 [17]. The stability of the substrate-bound *Tg*ASP5 structure was monitored using changes in RMSD, RMSF and radius of gyration with time (Fig. S2C, Fig. S2D, Fig. S2B). The substrate bound *Tg*ASP5 structure was stable compared to the apo structure as observed in the RMSD plot (Fig. S2C), but fluctuations were observed in the Insert-1 and Insert-2 regions as seen in the RMSF plot (Fig. S2C).

The Arg, present at P3 position of the TEXEL substrate gets accommodated in the S3 pocket (Fig. 3A, S7B) formed by the side chains of Glu506 and Gln549. The S3 pocket is strongly negatively charged (Fig. S6A). Similar feature has also been reported in the structure of *Pv*PMV. As both *Pv*PMV and *Tg*ASP5 recognise Arg at the P3 position of the substrate, so S3 pockets of these two enzymes are negatively charged, while in contrast S3 pockets of plasmepsin II is more positively charged [53]. Pepsin has a very small S3 pocket which favors the binding of alanine, while, renin, cathepsin D, cathepsin E all have preference for other hydrophobic residues at S3 binding pocket [54]. The negatively charged S3 binding pocket has also been found in other ASP5 enzymes- ASP5 *of Cyclospora cayetanensis)* (Fig. S6D), ASP5 of *Besnoitia besnoiti* (Fig. S6B), Aspartyl (Acid) protease of *Theileria orientalis* (Fig. S6C). The carboxylate group of Glu506 of the flap forms salt bridge interaction with the guanidine group of P3 Arg which is stable throughout the simulation process (Fig. 3G), it also forms polar interaction with the NE atom of the Arg side chain through OE1 and OE2 (Fig. 3A). The side chain of P3 Arg also has stable interaction with the carbonyl group of Gln549 sidechain in the S3 subsite (Fig. 3A, 3H). Thr686 present after the second catalytic motif (DSGT) also interacts with P3 Arg. The side chain hydroxyl group of Thr686 interacts with the main chain -NH group (Fig. 3A) as well as the carbonyl group of P3 Arg. These Glu506, Gln549, Thr686 are conserved among ASP5 enzymes (Fig. S3).

**Figure 3.**
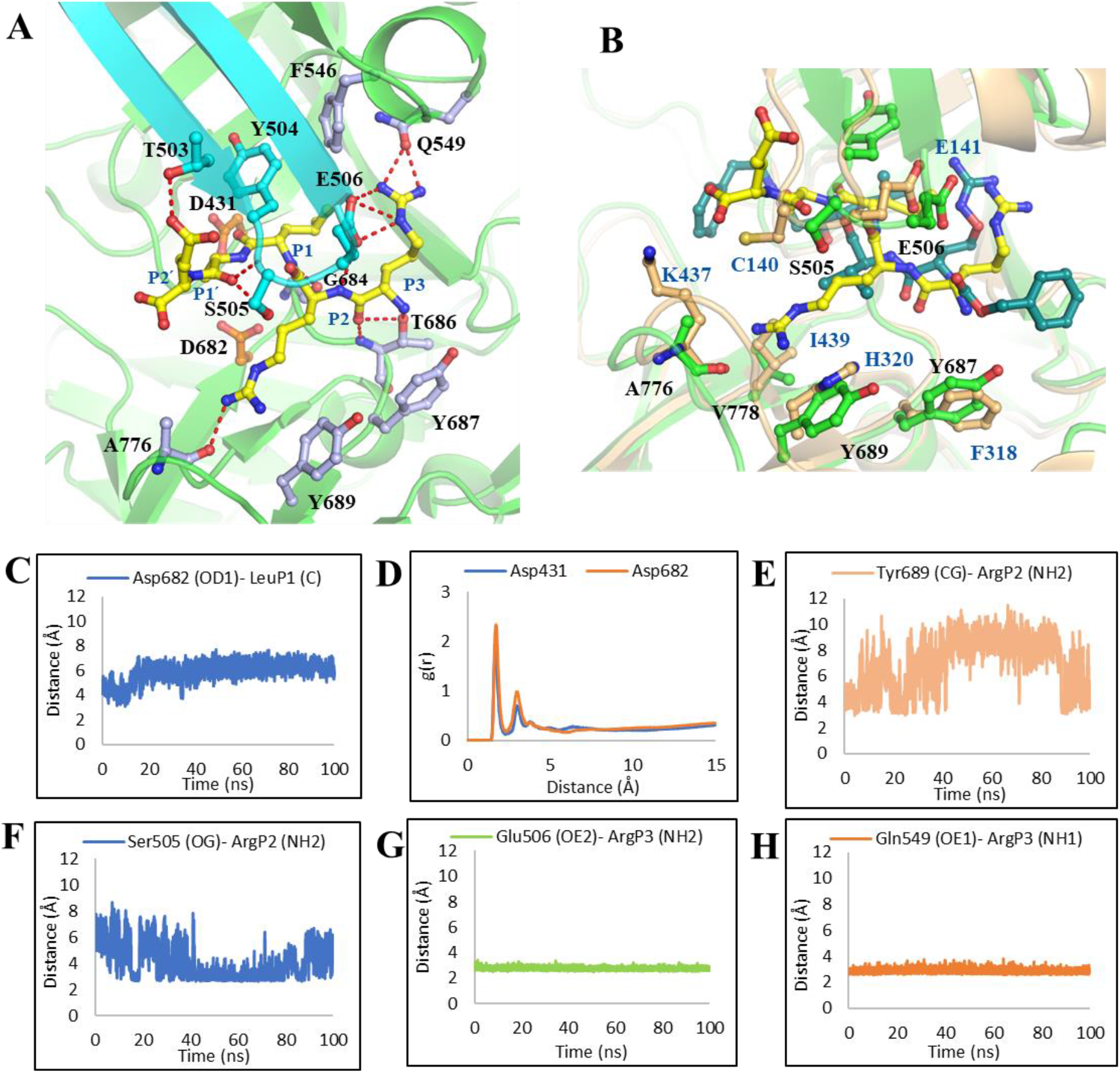
Interaction of TEXEL substrate in *Tg* ASP5 active site. **(A)** Binding mode of the TEXEL motif ^P3^RRL|AE^P2’^ in the active site pocket of *Tg*ASP5, the carbons of the peptide substrate are shown in yellow sticks. The two catalytic residues are shown in orange ball and sticks (carbon), other interacting residues are in light blue ball and sticks (carbon) flap is shown as cyan. The polar interactions are marked in red dashes. **(B)** Comparison of S2 binding pocket of *Pv*PMV and *Tg*ASP5. *Pv*PMV, *Tg*ASP5 both are represented in cartoon in light orange and green color, respectively, residues near S2 binding pocket are shown in ball and sticks. Carbon atoms of TEXEL substrate and PEXEL mimetic inhibitor WEHI-842 (4PK) are shown as sticks in yellow and deep teal, respectively. **(C)** The distance between the carbonyl carbon of P1 Leu of the substrate with OD1 atom of Asp682 during the simulation process. **(D)** The radial distribution function g(r) of the water molecule hydrogen atoms around the OD1 and OD2 of the Asp431 & Asp682 in the TEXEL-*Tg*ASP5 structure. The distance between **(E)** Tyr689 (CG)-ArgP2 (NH) of substrate and **(F)** Ser505 (OG)-ArgP2 (NH2) of substrate showing alternate interaction of P2 Arg with Tyr689 and flap Ser505. **(G)** Presence of stable salt bridge interaction between Glu506 (OE2) and NH2 atom of P3 Arg of substrate. **(H)** Hydrogen bonding interaction of Gln549 (OE1) with P3 Arg NH1 atom of substrate.

The Arg residue present at the P2 position of the substrate is well conserved among TEXEL substrates [17], unlike PEXEL of *Pf*PMV which usually has a small residue at this position [47]. The backbone N atom of the P2 Arg forms a stable hydrogen bonding interaction with the side chain carboxylate group of the flap Glu506 (Fig. 3A). The S2 binding pocket is mainly formed by Ser505, Ala776, Val778 and Tyr689 (Fig. S3B). The main chain carbonyl group of the Ala776 forms a hydrogen bonding interaction with the side chain of the P2 Arg (Fig. 3A). The Tyr689 forms a cation-π interaction with the guanidine group of the Arg (Fig. 3A). These two interactions were observed initially for around 0-40 ns and at around 90-100 ns (Fig. 3E, S7D); during 40-90 ns the side chain of P2 Arg was in constant interaction with Ser505 of flap tip (Fig. 3F). This suggests that the P2 Arg side chain is flexible and can interact with flap Ser505 or Ala776 and Tyr689, alternatively. The S2 pocket in *Pv*PMV has Lys437, His320, Ile439. (Fig. 3B), presence of His320 at the equivalent position of Tyr689 in *Tg*ASP5 might be the reason of absence of Arg at P2 position in PEXEL substrate. Besides that, the S2 pocket of *Tg*ASP5 also has Ser505 which is part of the flap region, at the same place Cys140 is present in *Pv*PMV which does not allow the binding of guanidium group of P2 Arg. The Ser505 is well conserved in ASP5 enzymes of other apicomplexan parasites, while all *Plasmodium* species contain Cys in the same position (Fig. S3). These key substitutions have made the *Tg*ASP5 active site to have requirement of Arg at P2 position of TEXEL substrate.

The S1 binding pocket that accommodates the side chain of Leu at P1 position of the TEXEL substrate is primarily hydrophobic, formed by flap Tyr504, Ile429, Phe546 and Ile554 (Fig. 3A, S7B, S7E), whereas the backbone of P1 Leu lies close to the N-terminal Asp431. The position of Ile429 and Ile554 (Fig. S3, S4) of *Tg*ASP5 are occupied by hydrophobic amino acids in other pepsin-like aspartic proteases. Apart from flap Tyr, vacuolar plasmepsins have another conserved Tyr residue in the S1 binding pocket, which is occupied by Phe546 in *Tg*ASP5 (Fig. S4) and is well conserved among ASP5 like enzymes (Fig. S3). The side chain of Ile429, Tyr504 and Phe546 helps the P1 Leu to remain stable in the S1 binding pocket through hydrophobic interaction (Fig. 3A, S7B). The nitrogen of the main chain of P1 Leu forms a hydrogen bonding interaction with main chain O atom of Gly684. The carbonyl carbon atom of the P1 Leu is at constant distance with the carboxylate group of catalytic Asp682 (Fig. 3C). Initially the distance was around 4Å which kept increasing gradually till it became 6Å after which the distance was constant. This distance is appropriate to accommodate a water molecule which is necessary for the nucleophilic attack on the carbonyl carbon of the P1 Leu. The carbonyl group of the scissile peptide bond points towards the Asp431. The radial distribution plot showed density of water molecules around Asp682 is more than Asp431 (Fig. 3D).

The P1́ Ala of the TEXEL substrate interacts with flap Ser505, the backbone O atom of P1́ Ala is present in hydrogen bonding distance with side chain OG atom and backbone N atom of the flap Ser505 (Fig. 3A, S7F). The S1́ binding pocket does not have a preference for any particular type of residue, all the identified *Tg*ASP5 substrates till now have either Ser, Ala or Asp in the P1́ site [17]. The same site can harbor polar, negatively charged and small neutral residues like Ala (Fig. S7A). In P2́ position, Glu is present for GRA16 and GRA21 which have a hydrogen bonding interaction with the side chain of Thr503 (Fig. S7G). This P2́ position is also not conserved among TEXEL substrates. The overall binding mode of the TEXEL substrate was found to be almost similar with the binding mode of PEXEL mimetic inhibitor WEHI-842 (4PK) present in the WEHI-842 bound crystal structure of *Pv*PMV (Fig. S7C).

### 3.5 Docking based screening of the inhibitors against *Tg*ASP5

As *Pf*PMV and *Tg*ASP5 share functional similarity therefore the PEXEL mimetic inhibitor WEHI-842 (4PK) was used to optimize the grid for docking studies using the later enzyme. The docking score for 4PK binding in *Tg*ASP5 active site (Fig. S8A) was found to be – 4.0 kcal/mol, and its docking pose is quite similar to the binding mode of the inhibitor observed in the complexed crystal structure (PDB ID: 4ZL4) of *Pv*PMV (Fig. S8B, S8C). Based on the grids optimized using 4PK, 80 peptidomimetic ligands of aspartic protease were obtained from Protein Data Bank (PDB) and screened as potential inhibitors. This set contains all peptidomimetic inhibitors of plasmepsins and a group of peptidomimetic BACE-1 and HIV-1 protease inhibitors. The docking-based screening suggests that BACE-1 inhibitors showed the highest affinity towards the *Tg*ASP5 binding site (Supplementary Table 1). Among the top ten hits, eight were found to be BACE-1 inhibitors. While for *Pf*PMV, HIV-1 protease inhibitors were the most effective inhibitors [49], reasons for the same could be some key substitutions in the S2 binding pocket of *Tg*ASP5. The substitutions include Arg773 for Val434 and Tyr689 for His320 in *Tg*ASP5 contribute to this behavior (Fig. S3). The substitution of Ser505 from Cys140 may also affect the inhibitor binding.

Both BACE-1 and *Tg*ASP5 are Golgi resident proteins and have similarities in active sites including Ala438 in the position of Trp. The flap motif is ^504^YSE^506^ in *Tg*ASP5 in place of ^132^YTQ^134^ of BACE-1 (Fig. S4). In place of the Glu506 of *Tg*ASP5, pepsin and *Pf*PMII has Thr/Ser respectively. Among the screened ligands, BACE-1 inhibitor QBH has the highest docking score of −6.73 kcal/mol. KNI-10333 (8VO), an inhibitor of vacuolar plasmepsins also showed good docking of −5.56 kcal/mol, but the orientation of the binding pose is exactly opposite to the crystal structure of *Pf*PMII-8VO bound complex (PDB ID: 5YIC) [55]. Presence of Tyr689 reduces the space to incorporate the phenylcyclopentane ring structure. 8VO can be well stabilized by forming interaction with catalytic Asp682 and other active site residues (Fig. S11D). A similar binding mode is also observed for KNI-10343.

### 3.6 Mode of Binding of a potent inhibitor RRL_Statine_ (WEHI-586) in *Tg*ASP5 active site

RRL_Statine_ contains the mimics of P3 Arg and P2 Arg of the TEXEL substrate along with the statin scaffold (Fig. 4A). It has phenyl groups at both terminals and a sulfonamide group beside the P3 Arg. The carbonyl group of the P1 Leu is replaced by the hydroxyl group of the statin. The central hydroxyl group makes constant hydrogen bonding interaction with C-terminal catalytic Asp682 (Fig. 4B, 4C). At the S3 binding pocket, Arg of this RRL_Statine_ forms salt bridge interaction with Glu506 similar to substrate. The Thr686 residue also forms polar interaction with the main chain of the Arg. The presence of sulfonamide group also stabilizes the interaction with Thr686 as O44 of the sulfonamide group also interacts with the OG1 atom of the Thr686. At the S2 binding pocket, the Arg side chain interacts with Ser505 and Tyr689 in a similar fashion as was observed for the substate. These interactions are very necessary for RRL_Statine_ as replacing this Arg with Val decreases the inhibition potency [17]. The main chain of P2 Arg has stable interaction with flap Glu506. Overall, the binding pose of RRL_Statine_ are almost similar with the substrate (Fig. 4D).

**Figure 4.**
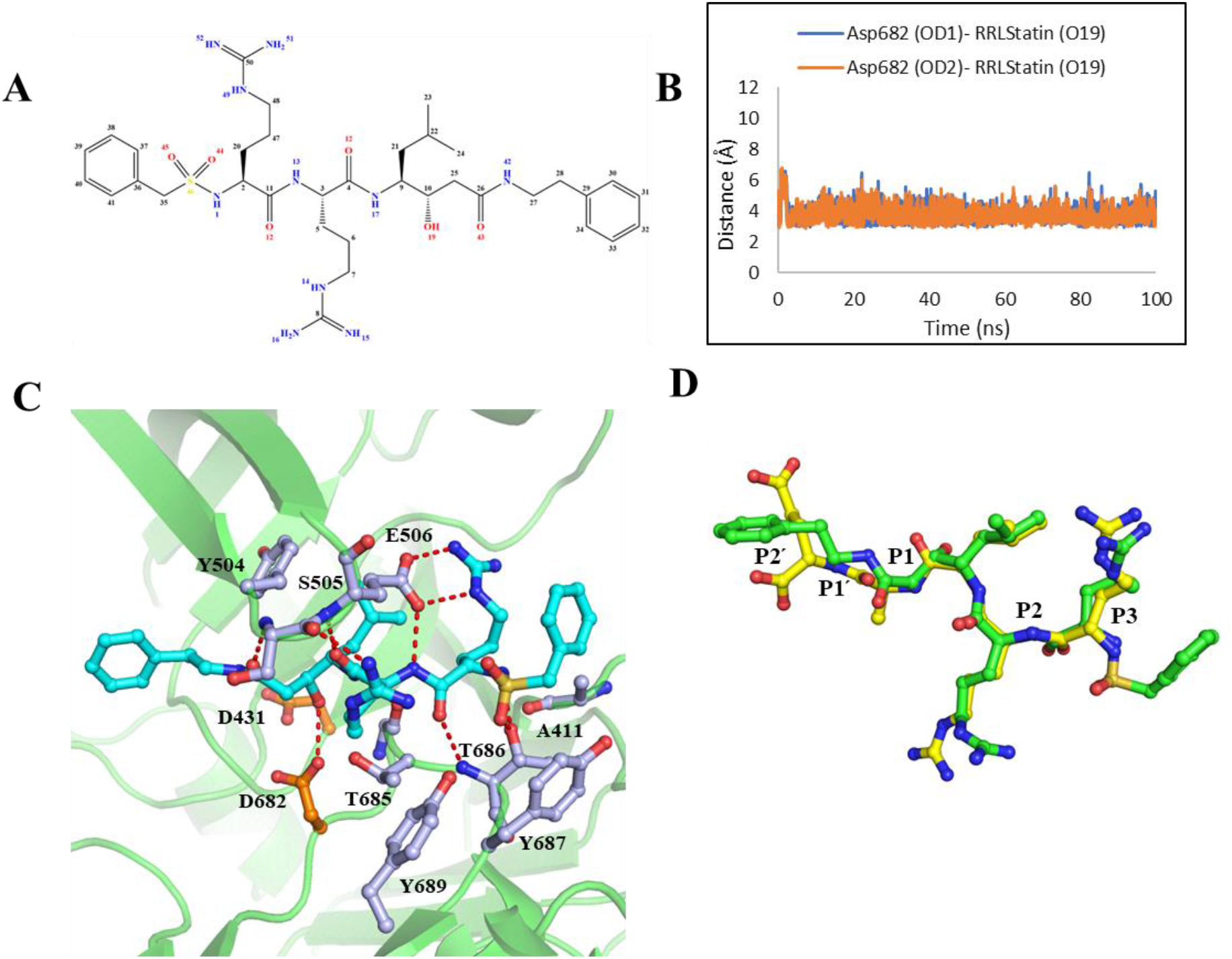
Molecular basis of inhibition of *Tg*ASP5 by the TEXEL mimetic inhibitor RRL_Statine_. **(A)** Covalent structure of RRL_Statine_. **(B)** Stable interaction between the central hydroxyl group (O19 atom) of statin moiety of RRL_Statine_ with OD1 and OD2 atom of the catalytic Asp682. **(C)** The interactions of RRL_Statine_ with active site residues. The H-bonding interactions formed by inhibitor (ball and sticks in cyan) in the active site of *Tg*ASP5 (green cartoon). The catalytic aspartates are indicted in orange ball and sticks while other active site residues are in light blue, polar interactions are marked in red dashes. **(D)** Alike binding mode of TEXEL substrate ^P3^RRLAE^P2’^ (yellow stick) and TEXEL mimetic RRL_Statine_ (green stick).

### 3.7 Simulation and MM-PBSA Binding Energy Calculation of ligand Bound Structures of *Tg*ASP5

Among the screened ligands, the nine best hits based on docking scores (Supplementary Table 1) were subjected to molecular dynamic simulations for 100 ns. After MD simulation the nine ligand-bound complexes along with 4PK and RRL_Statine_ bound *Tg*ASP5 were subjected to Molecular Mechanics Poisson-Boltzmann Surface Area (MM-PBSA) method to calculate the free energy of interaction of the *Tg*ASP5 with the ligands.

The binding energy for 4PK bound *Tg*ASP5 structure was found to be −96.5 kJ/mol. All the docked ligands selected for Simulations have better binding energy than 4PK. After MM-PBSA calculation, the binding energy of QBH was −278 kJ/mol, while that of SC6 was −321.2 kJ/mol (Fig. 5). Electrostatic energy has the major contribution to the higher binding energy for SC6. The binding energy of RRL_Statine_ was −72.3 kJ/mol, the inhibitor contributes to comparatively lesser electrostatic energy and relatively higher polar solvation energy than other screened ligands (Table-1).

**Table 1:** Energy value obtained for different ligands using MM-PBSA binding energy calculation

| Ligand | Van der Waals Energy (kJ/mol) | Electrostatic Energy (kJ/mol) | Polar Solvation Energy (kJ/mol) | SASA Energy (kJ/mol) | Binding Energy (kJ/mol) |
| --- | --- | --- | --- | --- | --- |
| SC6 | -280.9 ± 0.7 | -509.6 ± 0.8 | 499.0 ± 0.6 | -29.7 ± 0.0 | **-321.2 ± 0.8** |
| ZY1 | -288.3 ± 0.7 | -451.0 ± 1.0 | 484.6 ± 0.9 | -28.6 ± 0.0 | **-283.3 ± 0.7** |
| QBH | -220.8 ± 0.7 | -485.0 ± 0.9 | 453.8 ± 0.9 | -26.0 ± 0.0 | **-278.0 ± 0.8** |
| ZY0 | -262.2 ± 0.8 | -434.0 ± 1.5 | 491.5 ± 1.2 | -29.1 ± 0.0 | **-234.0 ± 0.9** |
| ZY4 | -219.8 ± 0.6 | -418.4 ± 1.0 | 432.1 ± 0.9 | -26.4 ± 0.0 | **-232.4 ± 0.8** |
| ZY2 | -229.6 ± 0.8 | -379.8 ± 0.9 | 426.1 ± 0.9 | -26.5 ± 0.0 | **-209.7 ± 0.8** |
| CM7 | -167.7 ± 0.6 | -304.6 ± 0.8 | 282.5 ± 0.6 | -19.7 ± 0.1 | **-209.6 ± 0.7** |
| AYH | -228.5 ± 0.5 | -78.0 ± 0.4 | 202.6 ± 0.5 | -24.8 ± 0.0 | **-128.7 ± 0.6** |
| 8VO | -260.3 ± 0.8 | -163.3 ± 0.9 | 350.4 ± 1.3 | -30.1 ± 0.1 | **-103.4 ± 0.8** |
| 4PK | -276.2 ± 0.6 | -100.0 ± 0.6 | 312.9 ± 1.0 | -33.2 ± 0.0 | **-96.5 ± 0.9** |
| RRL<sub>Statine</sub> | -256.3 ± 0.8 | -179.4 ± 1.3 | 394.6 ± 1.7 | -31.2 ± 0.1 | **-72.3 ± 1.0** |

**Figure 5.**
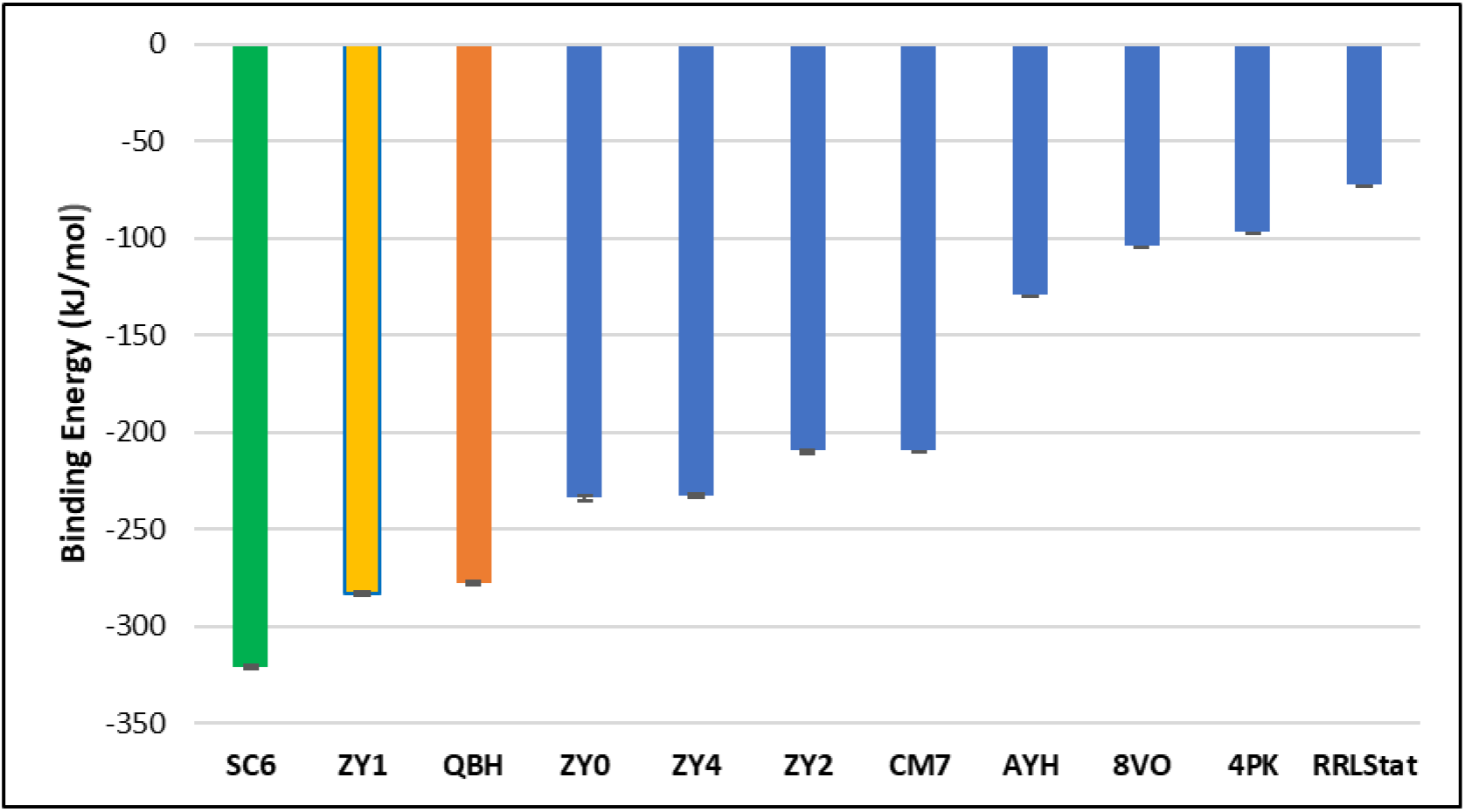
Comparison of ligand binding energies with *Tg*ASP5. Mean and standard deviation values of binding energy calculated by MM-PBSA method. The values are plotted along Y-axis and X-axis have different ligands.

### 3.8 Binding and Interaction of SC6 in *Tg*ASP5 Active Site

SC6 is a hydroxyethylamine scaffold-based aspartic protease inhibitor [56], the potential binding mode of the compound to *Tg*ASP5 was studied in detail as it shows the best binding energy using MM-PBSA approach [40]. The SC6 bound *Tg*ASP5 structure remained stable throughout the simulation observed by the change in radius of gyration, RMSD (Fig. S2B, S2C). The mode of binding of SC6 has similarity with the binding mode of substrate in P3 and P1 position and with the main chain at P2 position (Fig. 6A, Fig. 3A). The central hydroxyl group is placed at a similar position to the carbonyl group of the P1 Leu of the substrate. The -OH group of SC6 interacts with the catalytic Asp431 and the interaction is stable for 100 ns (Fig. 6B). Both the Asp431 and Asp682 forms interaction with the -NH group of the five-membered ring beside the central hydroxyl group (Fig. 6C, 6D). Presence of the - OH group and engagement of both catalytic Asp in interaction with ligand removes the tendency of water molecule to be present near them (Fig. 6G). Phe546 which is present in the active site forms π - π stacking interaction (Fig 6F) with the fluorine containing phenyl group of the inhibitor. Two fluorine atoms are present in SC6 also increases its potency by forming halogen interaction with the active site of *Tg*ASP5. The carbonyl oxygen of Leu545 forms an interaction with the fluorine of SC6 (Fig. 6F). The other fluorine atom forms interaction with the catalytic Asp431 (Fig. 6F).

**Figure 6.**
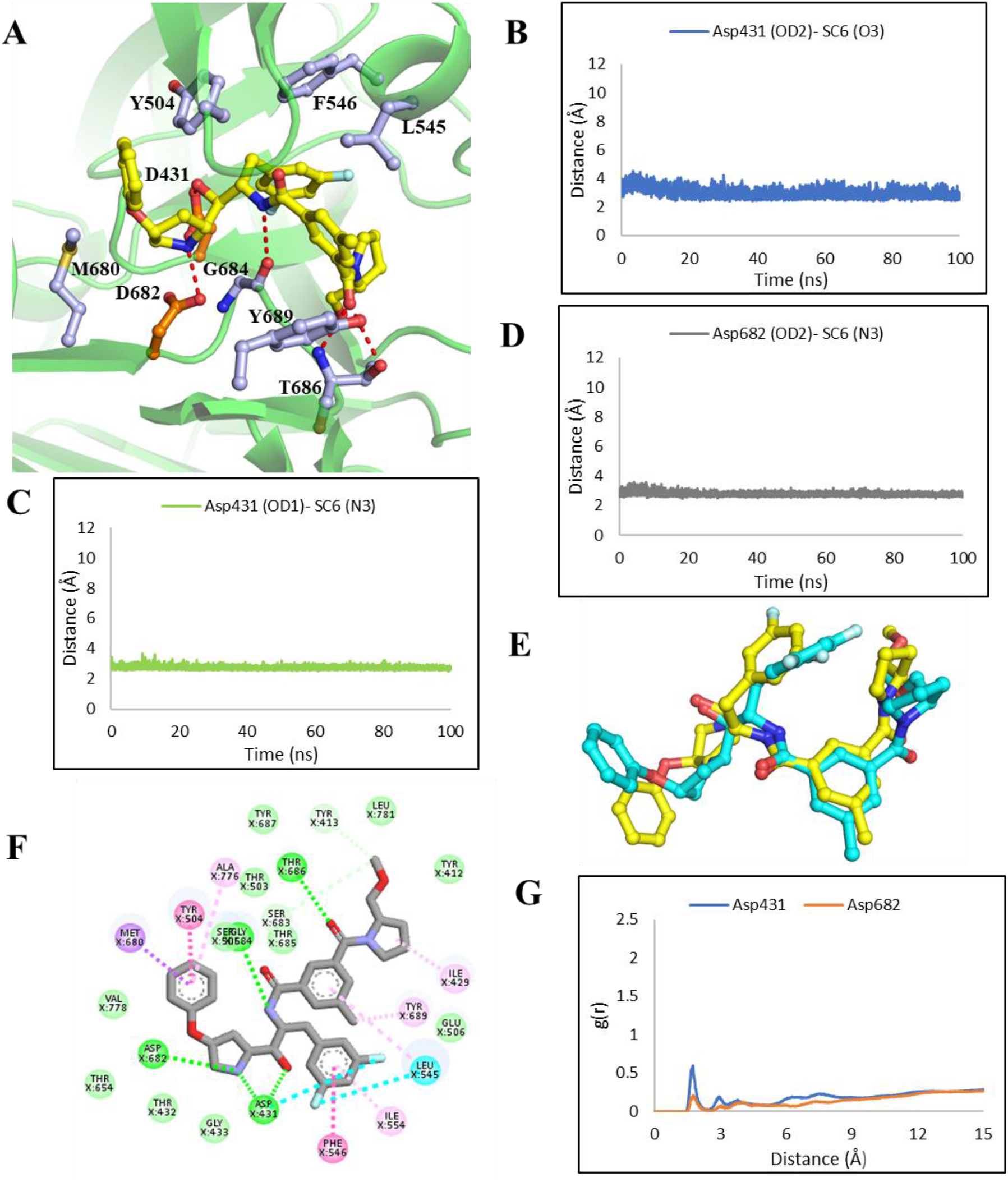
Binding mode of SC6 in active site pocket of *Tg* ASP5. **(A)** The zoomed-in view of the SC6 ligand binding to the active site pocket. The carbon atoms of SC6 are indicated in yellow ball and sticks, catalytic Asp431 & Asp682 as orange ball and sticks while other active site residues as light blue ball and sticks. The polar interactions are marked with red dashes. **(B)** The Asp431 forms stable interaction with O3 atom of SC6, which is part of the central hydroxyl group of the inhibitor through OD2 atom. **(C)** The other oxygen of carboxylate group OD1 forms hydrogen bond with backbone N3 atom of SC6. **D** The Asp682 also forms the stable hydrogen bond with the N3 atom of SC6. **(E)** The binding mode of SC6 is similar with the binding pose found in SC6-BACE-1 complex crystal structure (PDB ID: 2QMG). The binding pose of SC6 in *Tg*ASP5 active site is represented in yellow (carbon), for BACE-1, it is represented in cyan colored carbon. **(F)** The interaction diagram of SC6 with active site residues are shown in 2D representation. **(G)** The radial distribution function g(r) of the water molecule hydrogen atoms around the OD1 and OD2 of the Asp431 & Asp682 in the SC6-*Tg*ASP5 structure.

The Tyr504 of flap has an amide- π stacking interaction with SC6 (Fig. 6F). The main chain as well as side chain of Thr686 forms polar interaction with O1 atom of SC6 (Fig. 6A). A huge number of hydrophobic interactions is also observed in the SC6 bound structure of *Tg*ASP5 (Fig 6F). The overall binding mode of SC6 was almost similar what was observed in the crystal structure bound with BACE-1 (Fig. 6E).

Due to the strong binding of the SC6 with *Tg*ASP5, the movement core aspartic protease fold also gets reduced compared to the apo structure, which can be observed in the porcine plots based on the first and second principal components (Fig. S9). The substrate-protein complex also showed similar behavior (Fig. S9B, 9E). The movement of the flap region observed in the apo *Tg*ASP5 structure (Fig. S9A), became rigid after substrate or ligand binding.

### 3.9 Interaction with the other Inhibitors

Hydroxyethylamine-based transition state mimetic inhibitor ZY1 (PDB ID: 2WF1) also has a greater affinity for *Tg*ASP5 as MM-PBSA binding energy was −283.3 kJ/mol. The central hydroxyl group present between the P1 and P1́ position of the peptide substrate interacts with the catalytic Asp431 and Asp682 (Fig. S10A). The phenyl group is present under the flap makes π-π stacking interaction with Tyr504 of the flap and π-sigma interaction with Ile554. The methyl phenyl ether group of the ZY1 is present in the S1́ and S2́ binding pocket of *Tg*ASP5. The methyl group present close to the S2 binding pocket forms hydrophobic interaction with Met680, Ala 776, and Val778.

QBH is a cyclic hydroxyethylamine based inhibitor with cyclic sulfone [57]. The N-terminal catalytic Asp431 interacts with the -OH group of the inhibitor, which is the key interaction to inhibit the activity of the enzyme. It also interacts with the flap Ser505. The five-membered ring of the inhibitor which is present close to the P3 binding pocket makes hydrophobic interaction with Tyr412 and Tyr410. Lys550 forms polar interaction with the inhibitor from one end. The isopropyl phenyl group of QBH makes hydrophobic interactions Val778, Met680. (Fig. S10B).

A bicyclic heterocyclic biaryl hydroxyethylamine-based inhibitor ZY0 [58], have a binding energy of −234.02 kJ/mol. The central hydroxyl group of ZY0 interacts with Asp682 and Thr685 of the second catalytic motif. It also forms polar interaction with flap Ser505, Thr686 and Tyr687; and hydrophobic interaction with Tyr504, Tyr687, Ile429, Ile510 and Phe546 (Fig. S10C). ZY4, ZY2 both are tricyclic sulfonamide inhibitors [58,59] interacts with the catalytic Asp682 through hydrogen bonding (Fig. S10D) (Fig. S11A). ZY4 is an alpha-amino ketone-based inhibitor while ZY2 is the hydroxyethylamine based inhibitor.

Hydroxyethylamine-based inhibitor CM7 also binds to *Tg*ASP5 active site, where it places the tricyclic ring in the non-prime side of the active site, similar like BACE-1[60]. The hydroxyl group interacts with the N-terminal Asp while the neighboring amine group from ligand makes hydrogen bond with Asp682 (Fig. S11B). The BACE-inhibitor AYH, has a central hydroxyl group interacts with catalytic Asp682, Thr685 and flap Ser505 of *Tg*ASP5 (Fig. S11C).

## 4. Conclusions

*Tg*ASP5, an aspartic protease of *Toxoplasma gondii* plays a key role in host cell remodeling and is essential for the virulence as well as the fitness of the parasite. The enzyme is quite distinct from human aspartic proteases; it belongs to the different subfamily and has unique substrate cleavage site. The detailed structural studies of this enzyme have been carried out in this study; the enzyme has a pepsin-like aspartic protease fold with three inserts including A1B subfamily specific Nepenthesin I (Nep 1). Based on the analysis of binding mode of TEXEL substrate, Glu506, Gln549 and Thr686 residues play an important role in binding of P3 Arg. While S2 binding pocket specifically requires Arg at this position for *Tg*ASP5, which interacts alternately with flap Ser505 or Ala776 and Tyr689. This S2 subsite of *Tg*ASP5 is quite distinct from the *Pv*PMV which generally have smaller residues at this position. The Ser505, Ala776 and Tyr689 is necessary for catalysis of *Tg*ASP5. The only known inhibitor of *Tg*ASP5 is RRL_Statine_, its binding mode is similar to the substrate with central hydroxyl group containing non-cleavable amino acid statin which interacts with the catalytic Asp682. Based on the docking, simulation and binding energy calculation, few BACE-1 inhibitors including SC6, ZY1, QBH, containing the hydroxyethylamine-based scaffold were found to show better binding energy with the *Tg*ASP5 active site as compared to RRL_Statine_ and 4PK. Comparison of *Tg*ASP5 with BACE-1 shows certain variations in the S3, S2 and S2́ pocket of both the enzymes which can be exploited to design *Tg*ASP5 specific inhibitors. This study provides a comprehensive picture about the structural features and dynamics of *Tg*ASP5, along with molecular details of substrate binding pocket which will help to develop potent *Tg*ASP5 inhibitors.

---

**Figure S1.**
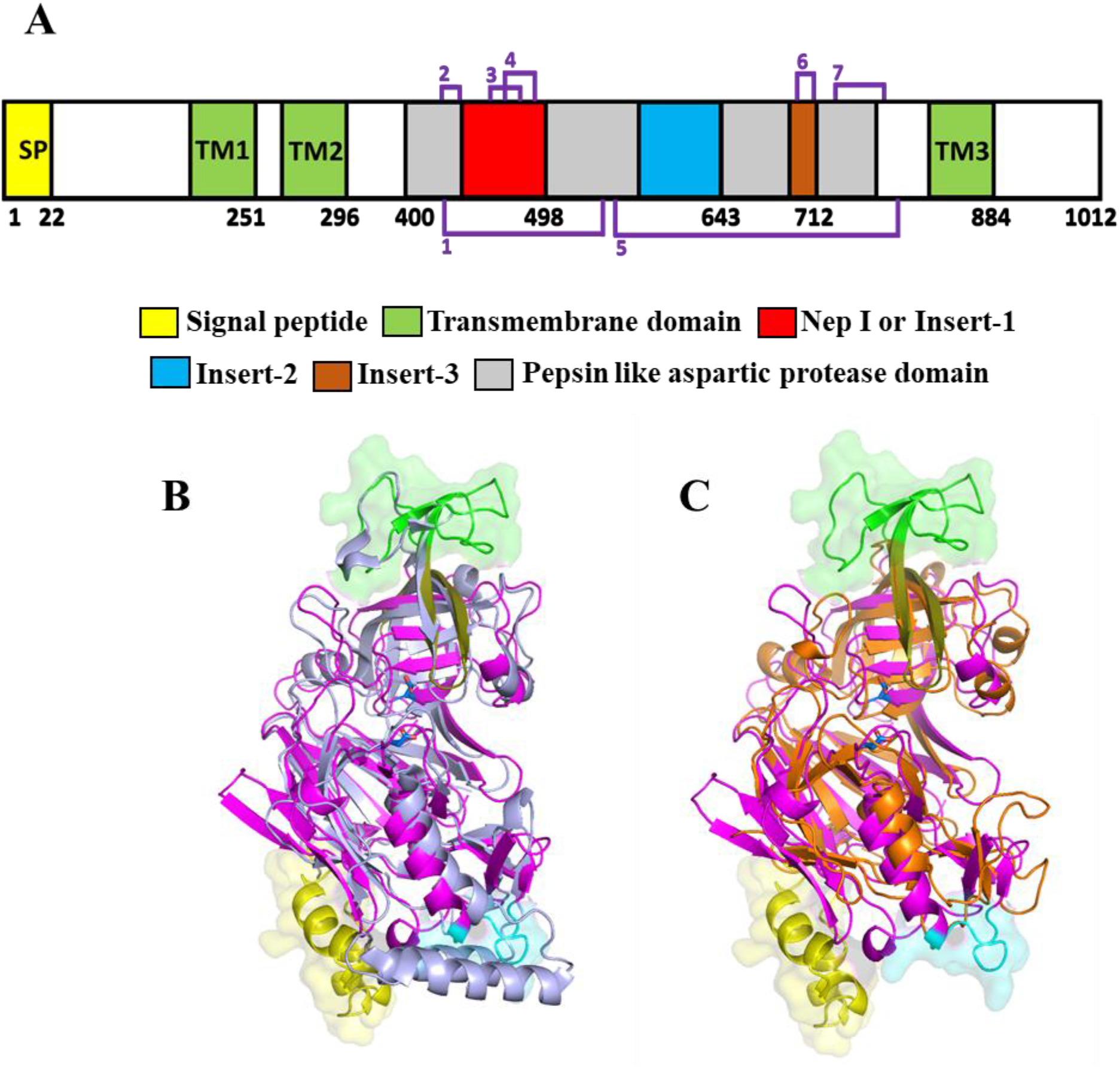
Domain organization of *Tg*ASP5. **(A)** The schematic representation of domain architecture of *Tg*ASP5. *Tg*ASP5 contains a Signal peptide and three Transmembrane domains (TM). The protease domain has pepsin-like aspartic protease fold along with three inserts. The Signal peptide and Transmembrane domains are marked with yellow and green color. The Insert-1, Insert-2, Insert-3 are shown in red, blue and brown color respectively and pepsin-like aspartic protease fold is represented in grey. The presence of seven disulphide bonds in the protease domain are shown in magenta. Structural comparison of *Tg*ASP5 with **(B)** *Pv*PMV (PDB ID: 4ZL4) and **(C)** *Pf*PMII (PDB ID: 1LF3). The structure of mature *Tg*ASP5 is represented as cartoon. The classical pepsin-like aspartic protease fold is shown in magenta and flap in deep olive, Insert-1 (Nep 1), Insert-2 and Insert-3 are shown in semi-transparent surface in green, yellow and cyan color. The *Pv*PMV is represented in light blue cartoon. *Pf*PMII structure is shown in orange cartoon.

**Figure S2.**
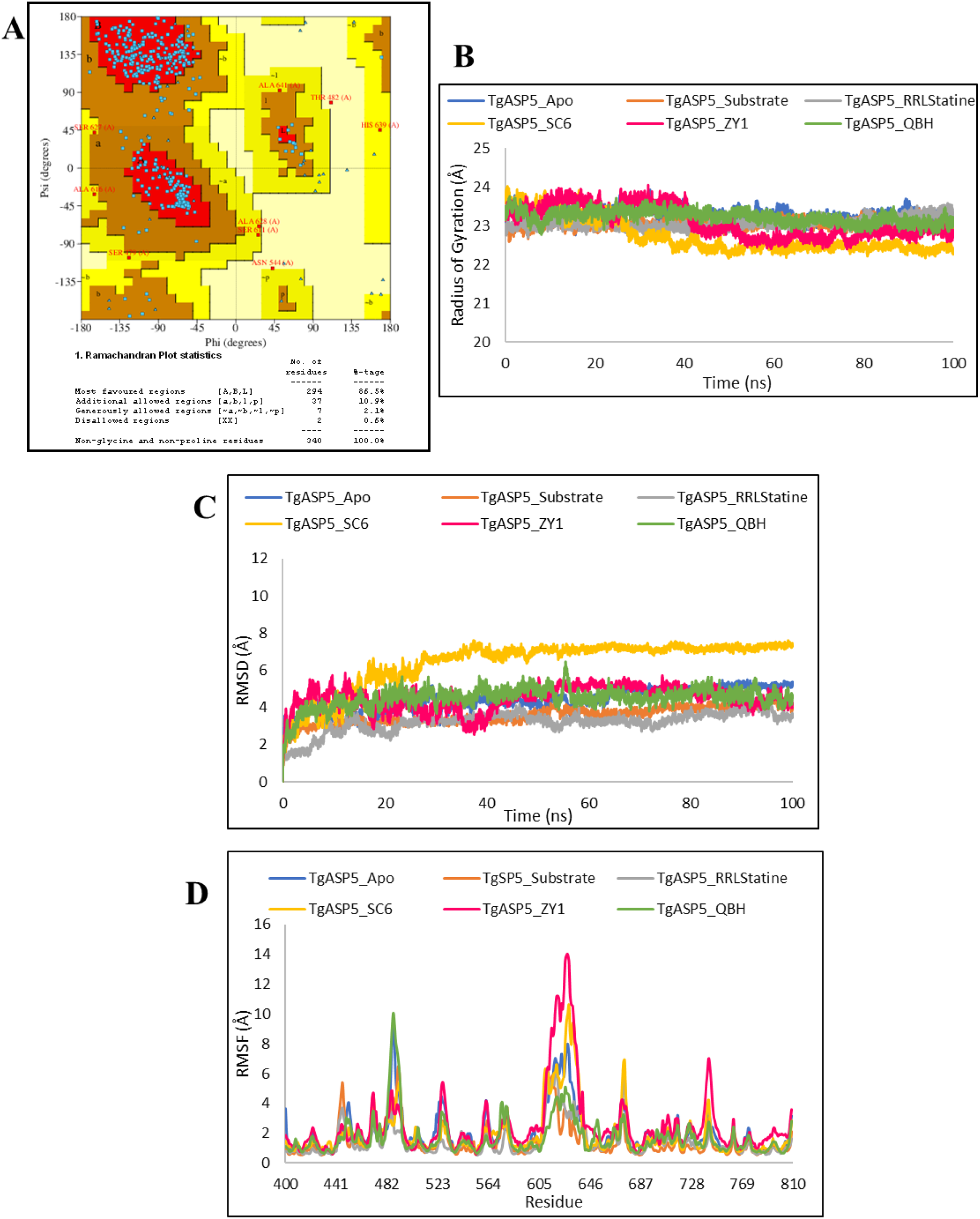
Validations of *Tg*ASP5 model and structural stability during simulations. **(A)** Ramachandran plot of initial model of *Tg*ASP5. The stability of the overall structure of Apo *Tg*ASP5, TEXEL substrate-*Tg*ASP5, ligand (RRL_Statine_, SC6, ZY1, QBH)-*Tg*ASP5 complexes were assessed with the help of **(B)**Radius of gyration (Rg) plot, **(C)** Root mean square deviation (RMSD) plot based on backbone and **(D)** Root mean square fluctuation (RMSF) of Cα atoms.

**Figure S3.**
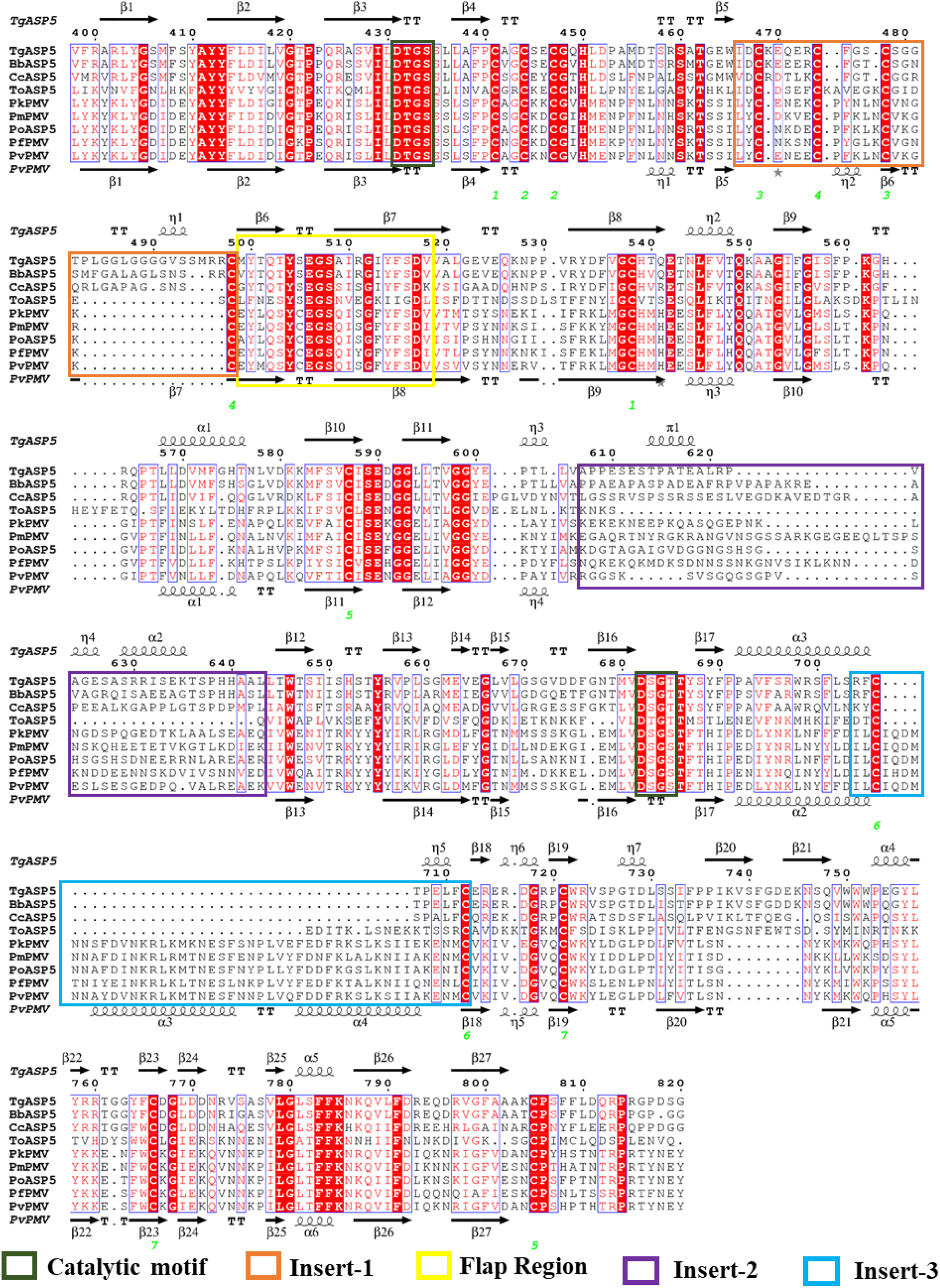
Comparison of sequences of different ASP5 enzymes. DTGS/DSGT motif, and flap region are marked as dark green and yellow boxes, Insert-1, Insert-2, Insert-3 are marked as orange and purple and cyan boxes respectively. Position of the disulphide bond is shown by green numbering below the alignment. Secondary structural elements of *Tg*ASP5 are shown on top and of *Pv*PMV (PDB ID: 4ZL4) is shown on bottom. The identical residues are shown in red background and similar residues in red color, generated by ESPript (https://espript.ibcp.fr/ESPript/ESPript/).

**Figure S4.**
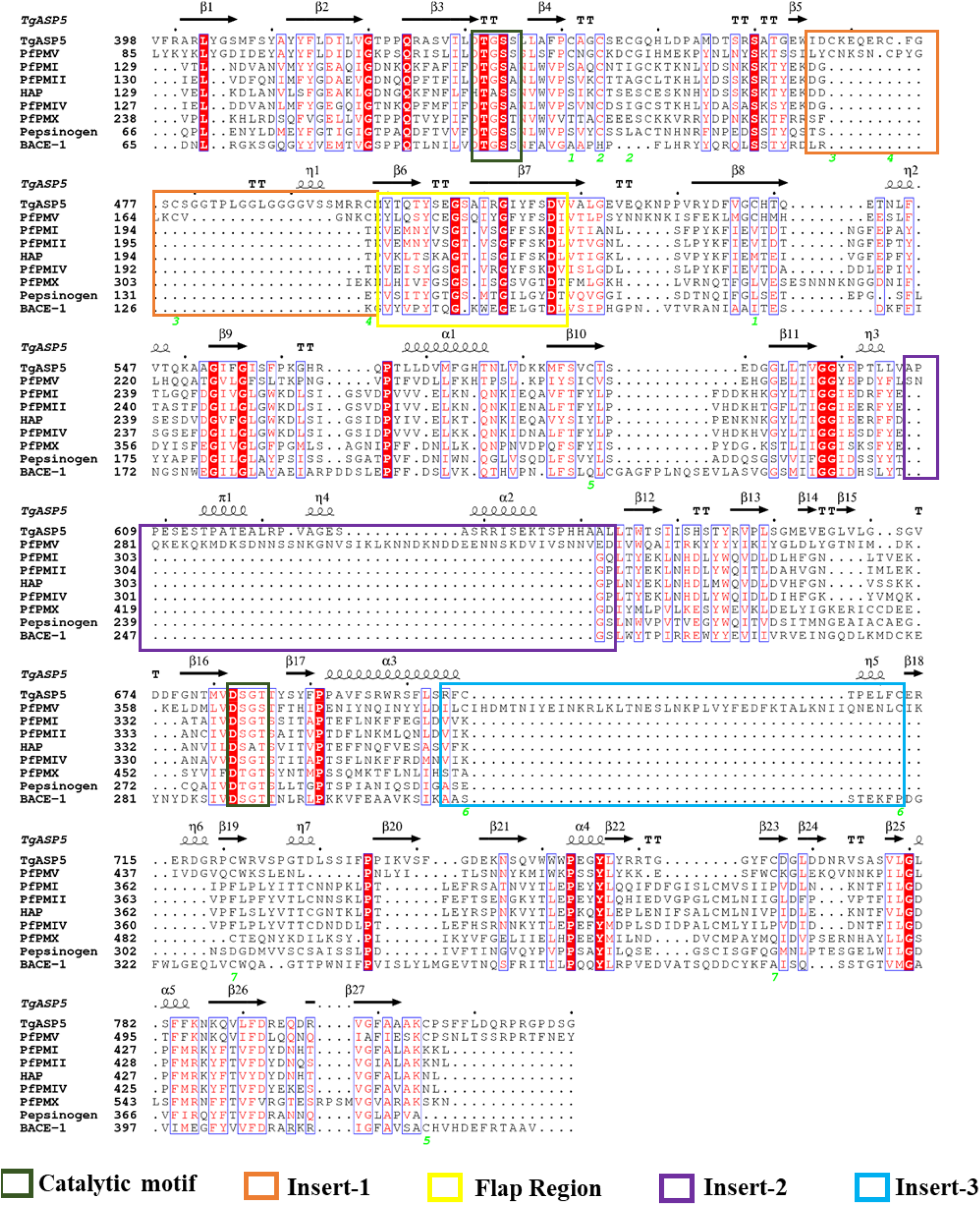
Sequence comparison of *Tg*ASP5 with pepsin and pepsin like aspartic proteases. DTGS/DSGT motif, and flap region are marked as dark green and yellow boxes, Insert-1, Insert-2, Insert-3 are marked as orange and purple and cyan boxes respectively. Position of the disulphide bond is shown by numbering below the sequences. Structural elements of *Tg*ASP5 are shown on top.

**Figure S5.**
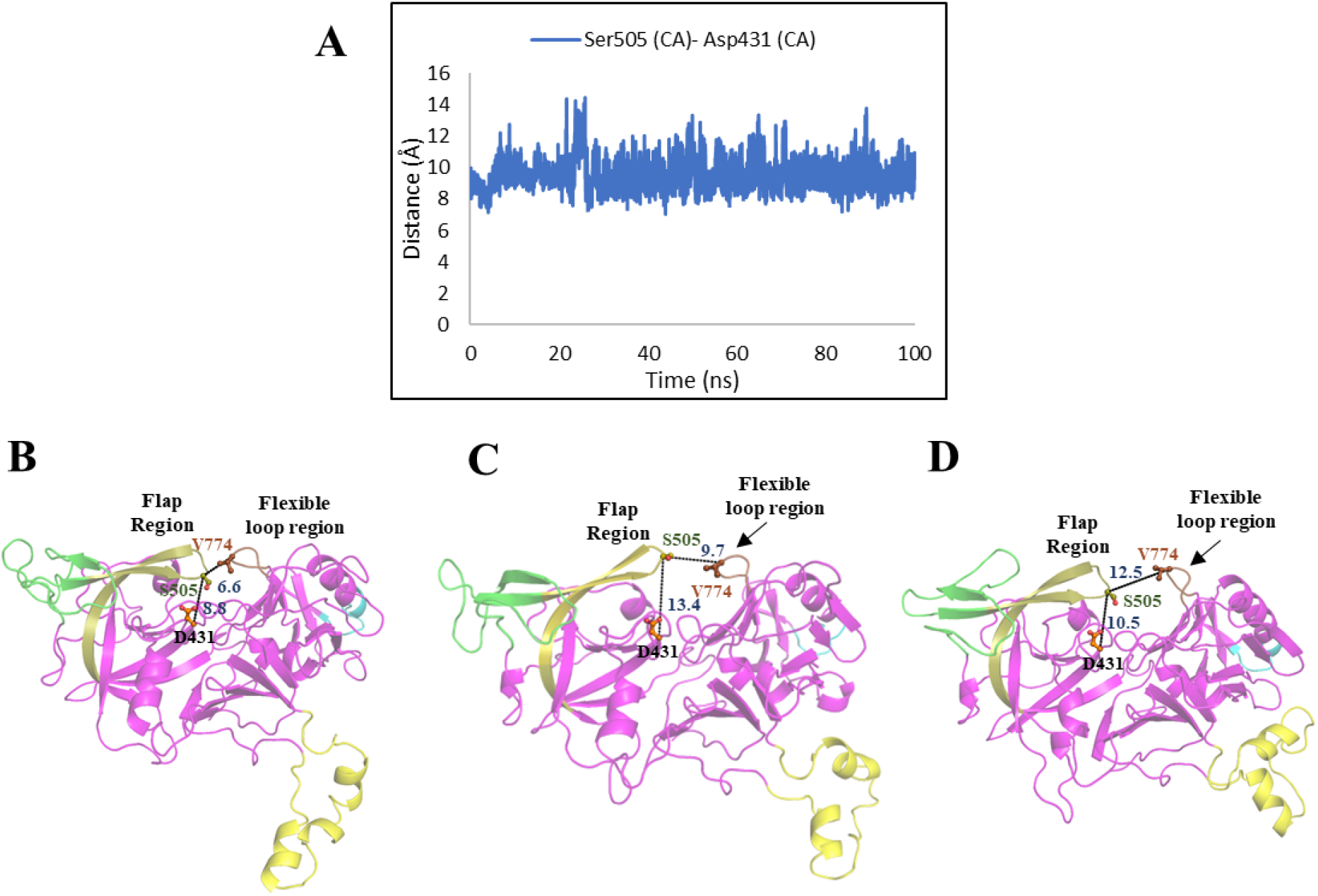
Dynamics of the flap in *Tg* ASP5 apo-structure. **(A)** Distance between the CA atom of flap tip Ser505 and the CA atom of N-terminal catalytic Asp431. **(B)** Close structure of active site pocket of *Tg*ASP5 at starting 4ns of the simulation. **(C)** Opening of active site pocket of *Tg*ASP5 structure at 26 ns. **(D)** *Tg*ASP5 active site pocket at 52 ns of MD simulation. The structure of apo *Tg*ASP5 is represented as cartoon. The classical pepsin-like aspartic protease fold is shown in magenta and flap in deep olive, flexible loop region in brown, while Insert-1, Insert-2 and Insert-3 are shown in green, yellow and cyan color. The opening and closing of the active site is happening with the dynamics of flap and the flexible loop region.

**Figure S6.**
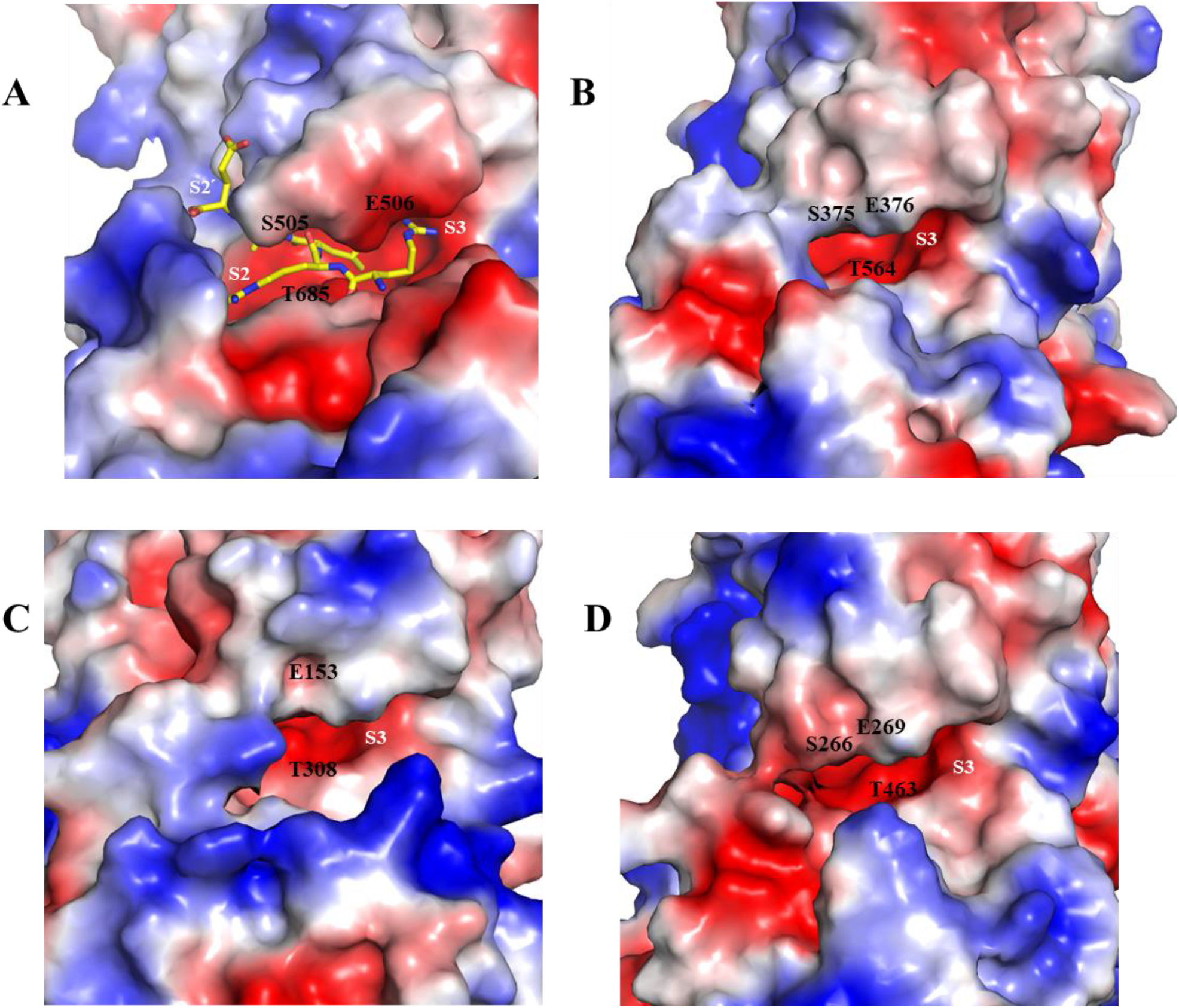
Substrate binding pocket of ASP5 enzymes. Presence of negatively charged S3 binding pocket in **(A)** *Toxoplasma gondii*, **(B)** *Besnoitia besnoiti*, **(C)** *Theileria orientalis* **(D)** *Cyclospora cayetanensis*. Electrostatic surface (Red: negative, blue: positive charge, white: neutral) representation of substrate binding pocket. The S3 sub sites are marked with magenta. The TEXEL substrate is represented in yellow sticks.

**Figure S7.**
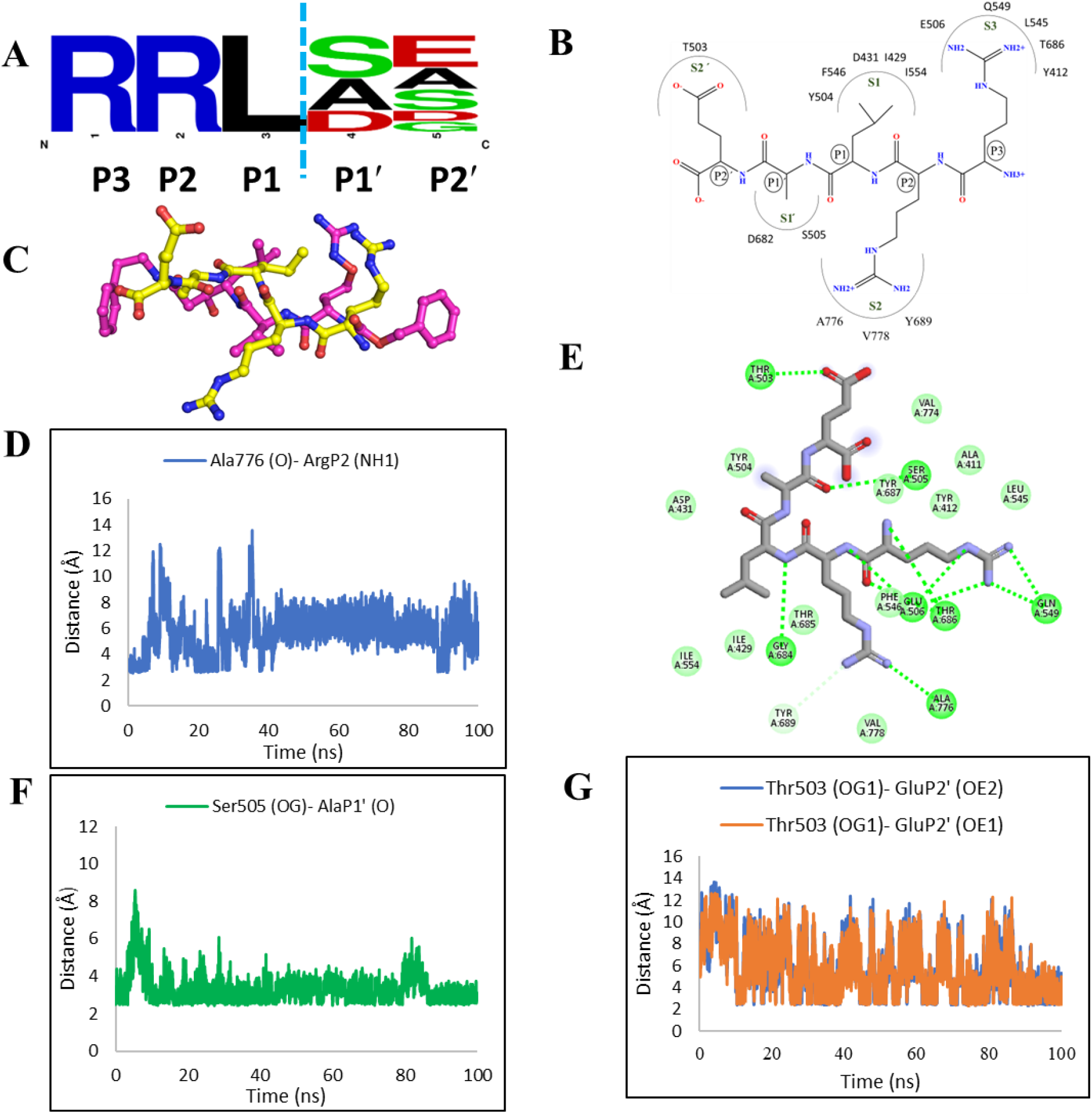
Molecular basis of recognition of TEXEL motif by *Tg*ASP5. **(A)** The variation of residues at P1` and P2` position of TEXEL substrate. The WebLogo (http://weblogo.berkeley.edu) is generated using the alignment of the known TEXEL substrate. The cyan dashed line is showing the cleavage site of TEXEL. **(B)** Schematic representation of binding of TEXEL substrates in *Tg*ASP5 active site. **(C)** Similarity in the predicted binding mode of TEXEL substrate with the PEXEL mimetic inhibitor 4PK as observed in the crystal structure (PDB ID: 4ZL4). The carbon atoms of TEXEL substrate and 4PK are shown in yellow and pink respectively. **(D)** Distance between backbone O atom of Ala776 with NH1 atom of P2 Arg. **(E)** 2D interaction diagram of TEXEL substrate-*Tg*ASP5 complex. **(F)** Distance between the backbone N atom of P1` Ala with backbone N atom and sidechain OG atom of Ser505 and **(G)**Sidechain OG1 atom of flap Thr503 with OE2 and OE1 atom of the carboxylate group of P2` Glu respectively.

**Figure S8.**
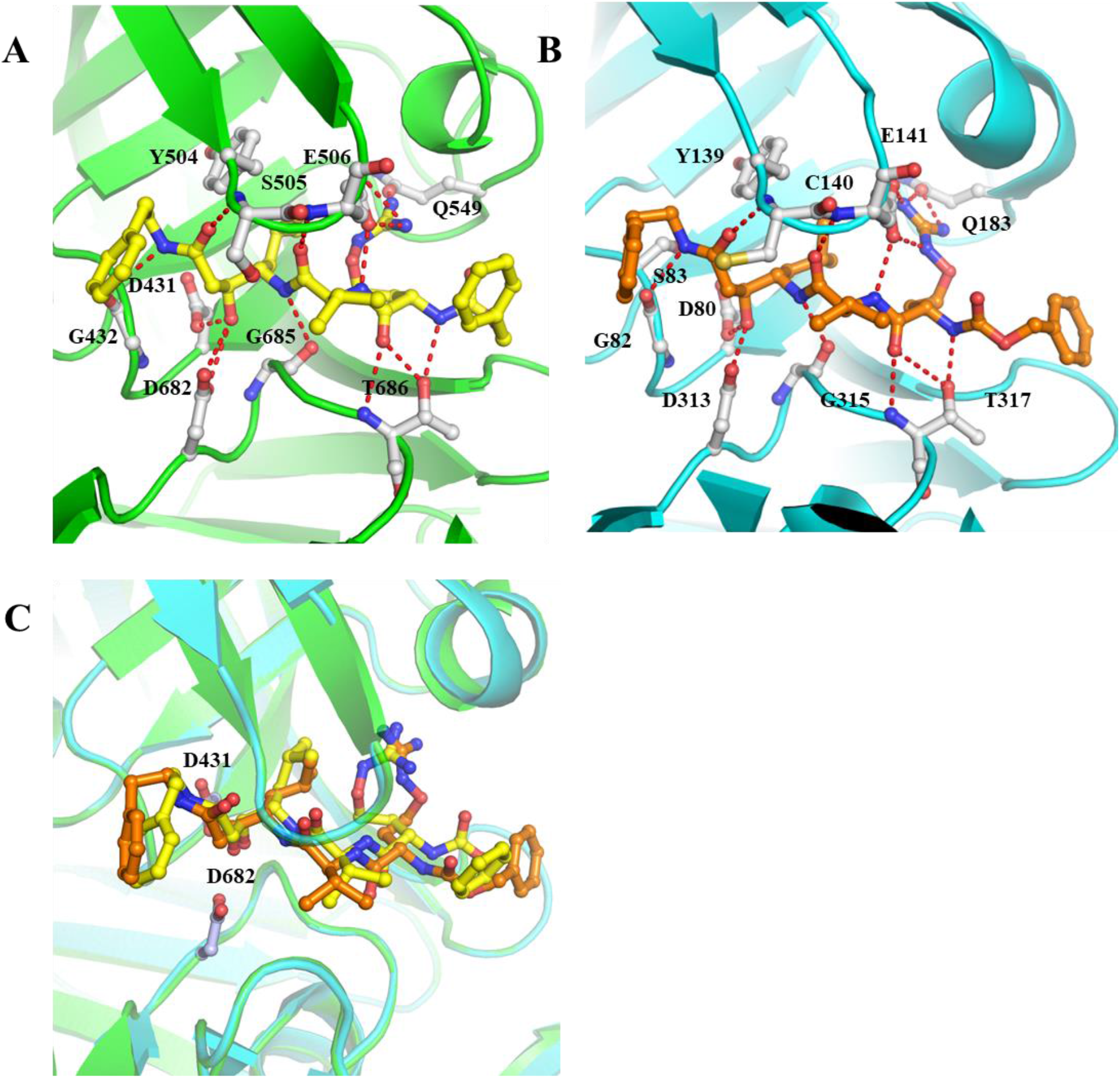
Docking of PEXEL mimetic inhibitor 4PK in *Tg*ASP5 active site. **(A)** The docked structure of *Tg*ASP5-4PK complex. The 4PK inhibitor is represented in yellow (carbon) ball and sticks. The interacting residues are represented in gray ball and sticks. The hydrogen bonds are marked with red dashed line. **(B)** The zoomed in view of *Pv*PMV-4PK complex crystal structure. The inhibitor is shown in orange ball and sticks. The active site residues are shown in gray ball and sticks. The hydrogen bonds are represented in red dashed line. The *Pv*PMV is represented in cartoon in cyan. **(C)** The comparison of the docked pose of 4PK with pose obtained in the crystal structure with *Pv*PMV. The docked pose of 4PK in *Tg*ASP5 active site is shown in yellow, while 4PK of the *Pv*PMV-4PK complex is shown in orange. The catalytic Asp431, Asp682 of *Tg*ASP5 are shown in gray colored carbon. *Pv*PMV is shown in cyan cartoon and *Tg*ASP5 in green cartoon.

**Figure S9.**
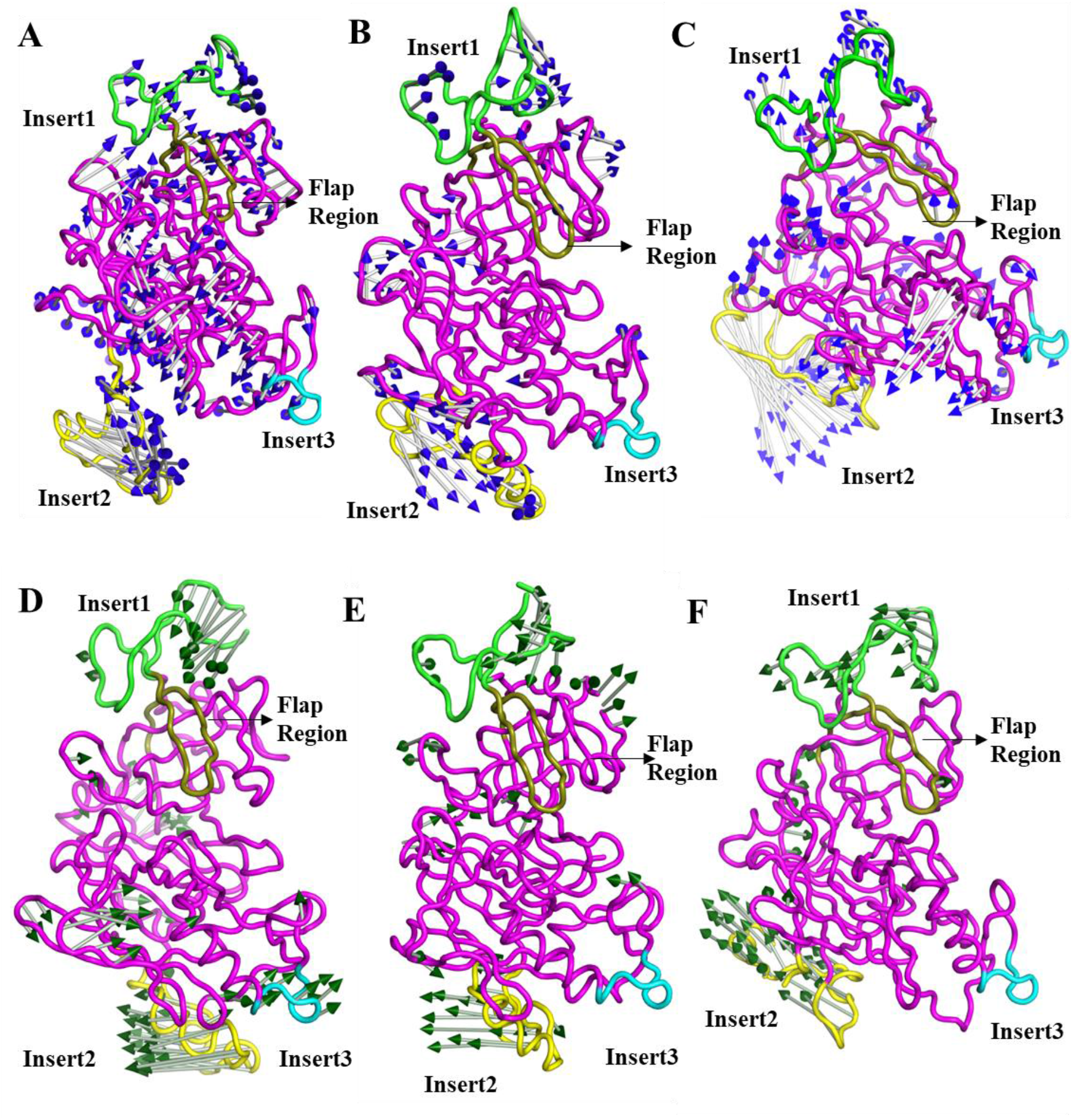
Porcine plots corresponding to PC1 & PC2 generated by Principal Component Analysis on MD trajectories of Apo, substrate and ligand bound *Tg*ASP5 structures. The first three porcine plots **(A) (B) (C)** corresponds to PC1 of Apo, TEXEL substrate-*Tg*ASP5, Ligand (SC6)- *Tg*ASP5 complexes respectively for 100 ns classical MD simulation. The blue arrows are showing the vector movement during the simulation process. The pepsin-like aspartic protease fold of *Tg*ASP5 is shown in magenta along with flap in deep olive, insert-1, insert-2, insert-3 are represented in green, yellow and cyan respectively. **(D), (E), (F)** corresponds to PC2 of apo, TEXEL bound and SC6 bound *Tg*ASP5 complexes, the movement is shown with green arrow.

**Figure S10.**
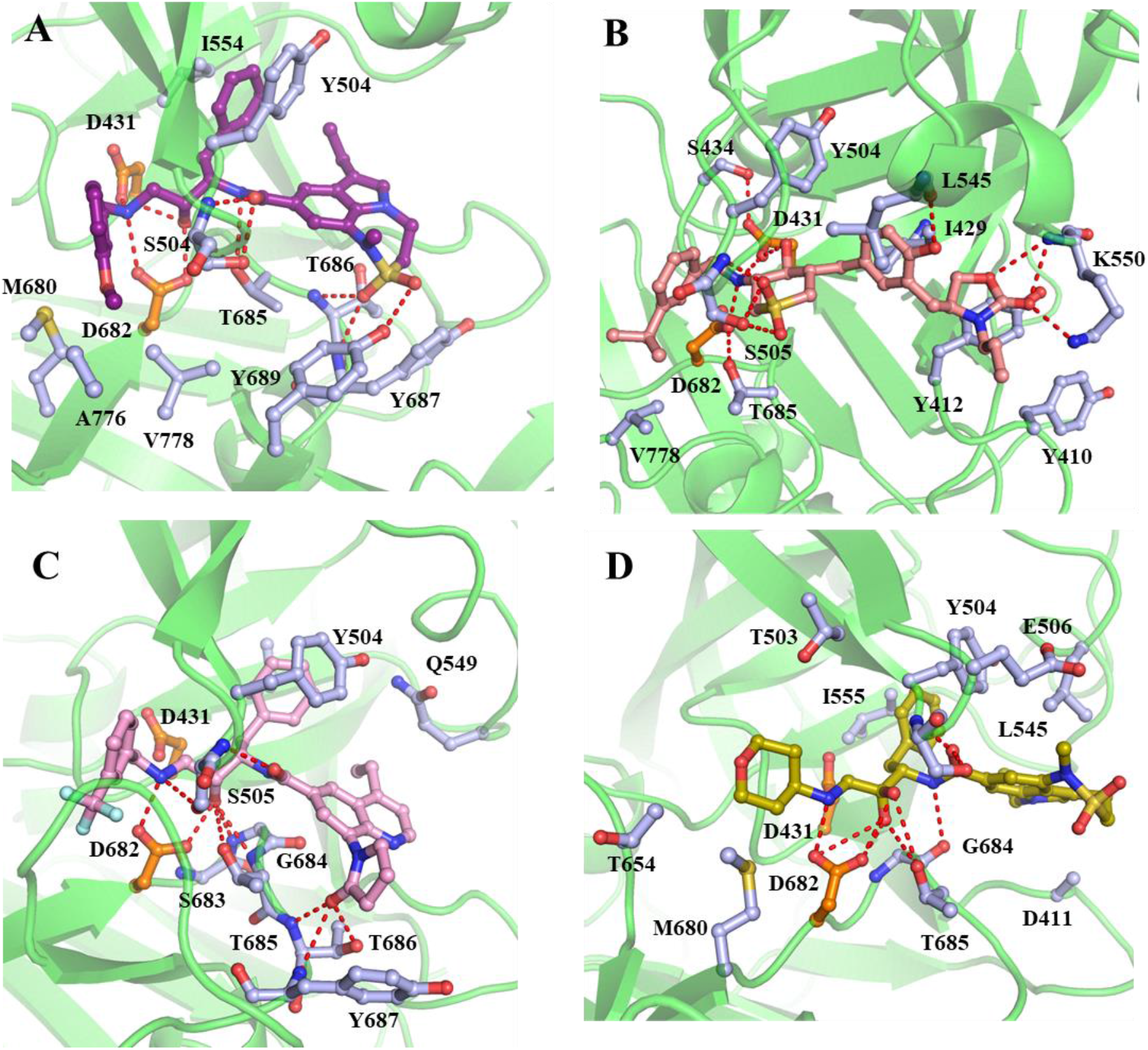
Interactions of ZY1, QBH, ZY0 and ZY4 with *Tg*ASP5. **(A)** Zoomed in view of active site of ZY1- *Tg*ASP5 complex. The bound inhibitor is shown in ball and stick model with carbon colored in purple. The residues involved in interactions are also shown in ball and sticks with carbon colored light blue, the catalytic Asp431 and Asp682 are shown in orange color carbon. **(B)** Interactions formed by QBH (wheat carbon) with the active site residues of *Tg*ASP5. Both the inhibitor and the residues involved in interactions are represented in ball and stick model, the catalytic Asp431, Asp682 are indicated in orange carbon, others are marked with light blue carbon. The polar interactions are marked with red dashed line. **(C)** The inhibitor ZY0 is represented in pink (carbon) ball and sticks, the catalytic Asp are indicted in orange (carbon) and other interacting residues in light blue ball and sticks. The hydrogen bonds are shown in red dashed line. **(D)** Zoomed in view of the *Tg*ASP5 active site bound with ZY4. The bound inhibitor is shown in olive (carbon) ball and sticks, the catalytic Asp are indicted in orange colored carbon and other interacting residues in light blue ball and sticks. The polar interactions are shown in red dashed lines.

**Figure S11.**
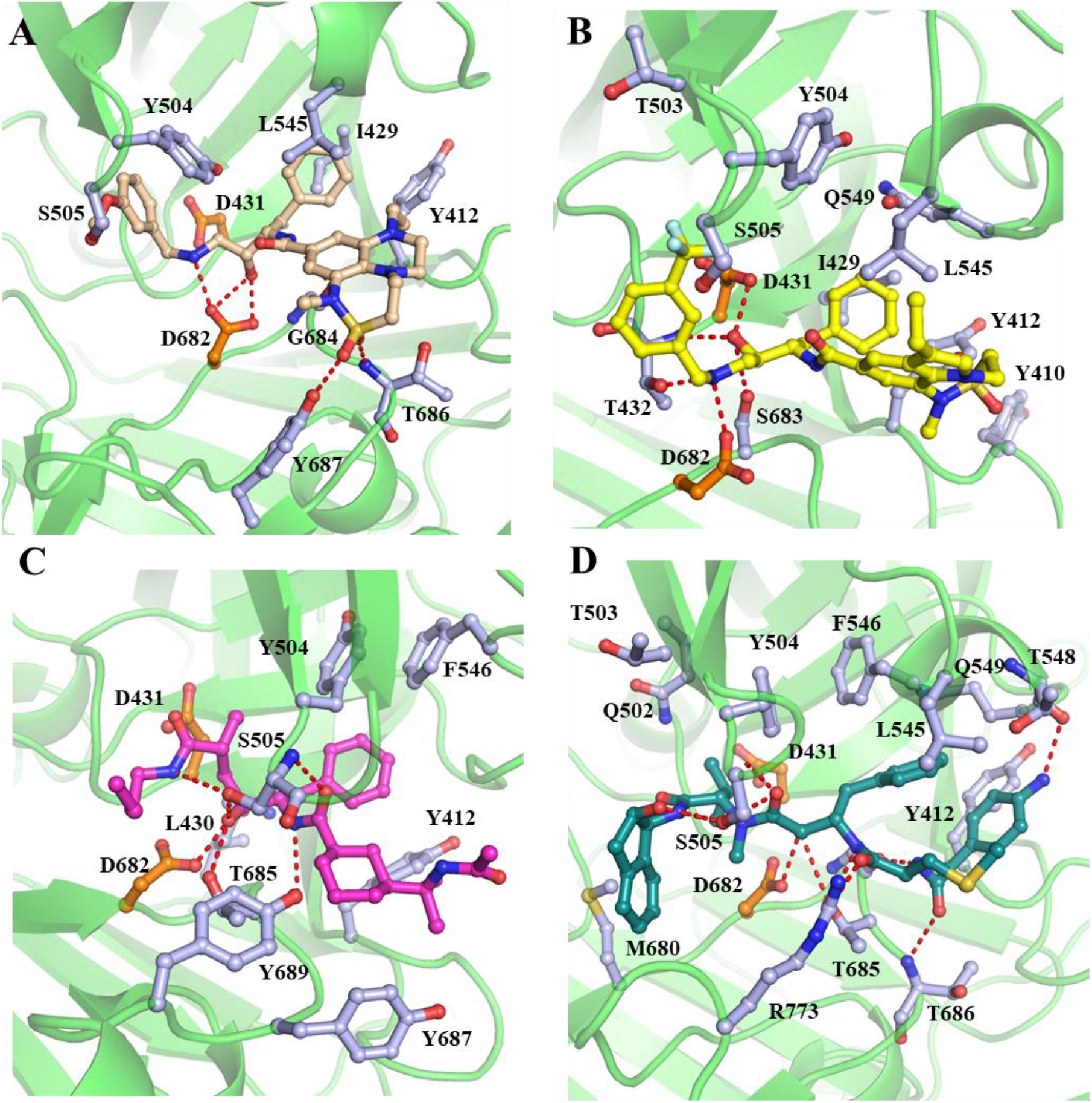
Active site of TgASP5-ZY2, TgASP5-CM7, TgASP5-AYH and TgASP5-KNI10333 complexes. **(A)** The inhibitor ZY2 is represented in wheat (carbon) ball and sticks, the catalytic Asp are indicted in orange (carbon) and other interacting residues in light blue ball and sticks. The hydrogen bonds are shown in red dashed line. **(B)** CM7 is represented in yellow (carbon) ball and sticks, the catalytic Asp are indicted in orange (carbon) and other interacting residues in light blue ball and sticks. The hydrogen bonds are shown in red dashed line. **(C)** Zoomed in view of the *Tg*ASP5 active site. The bound AYH inhibitor is shown in magenta colored carbon, the catalytic Asp are indicted in orange colored carbon and other interacting residues in light blue. The polar interactions are represented in red dashed line. **(D)** The inhibitor KNI1033 (8VO) is represented in deepteal (carbon) ball and sticks, the catalytic Asp are indicted in orange (carbon) and other interacting residues in light blue ball and sticks. The hydrogen bonds are shown in red dashed line.

**Supplementary Table 1:**
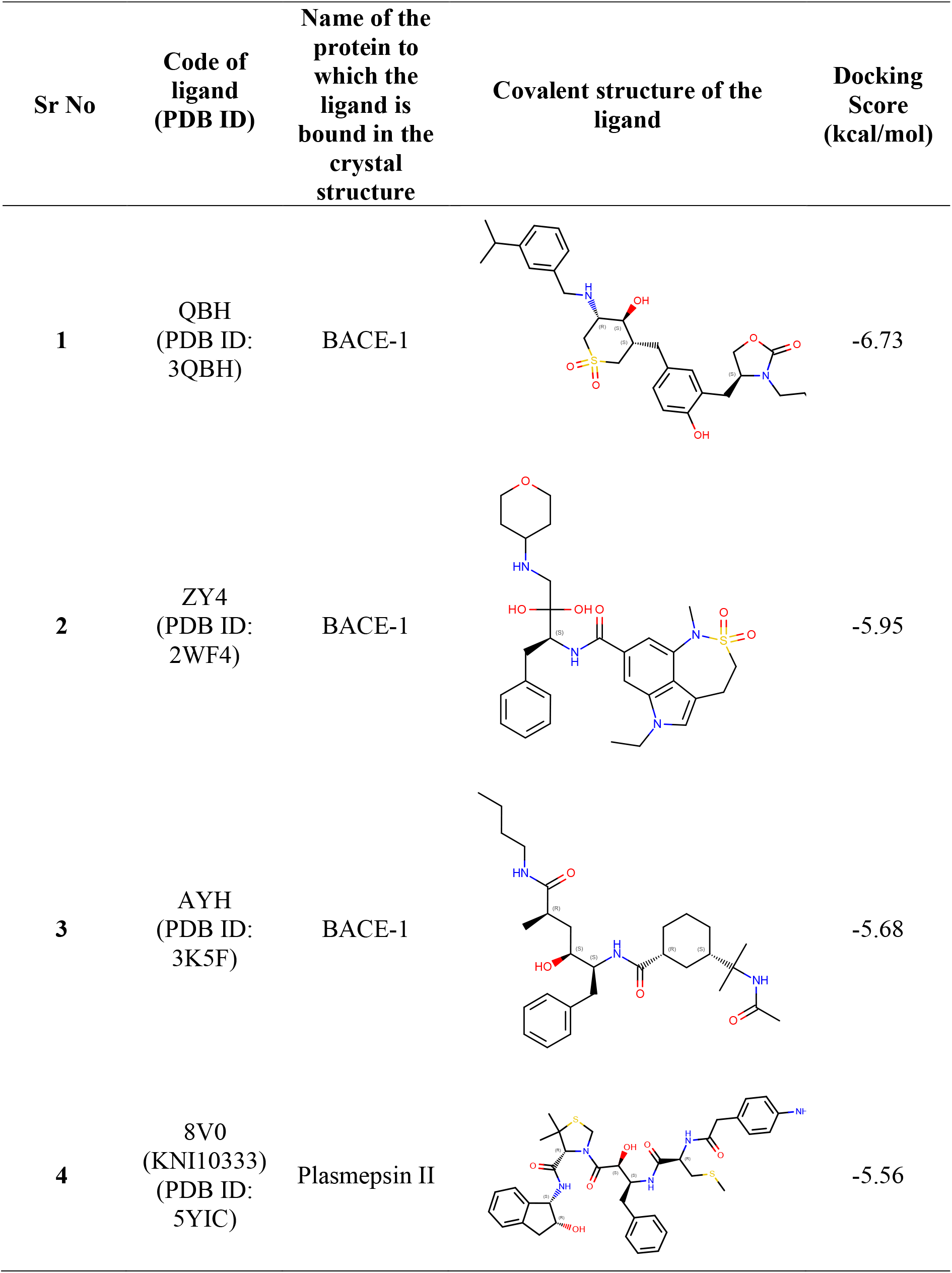
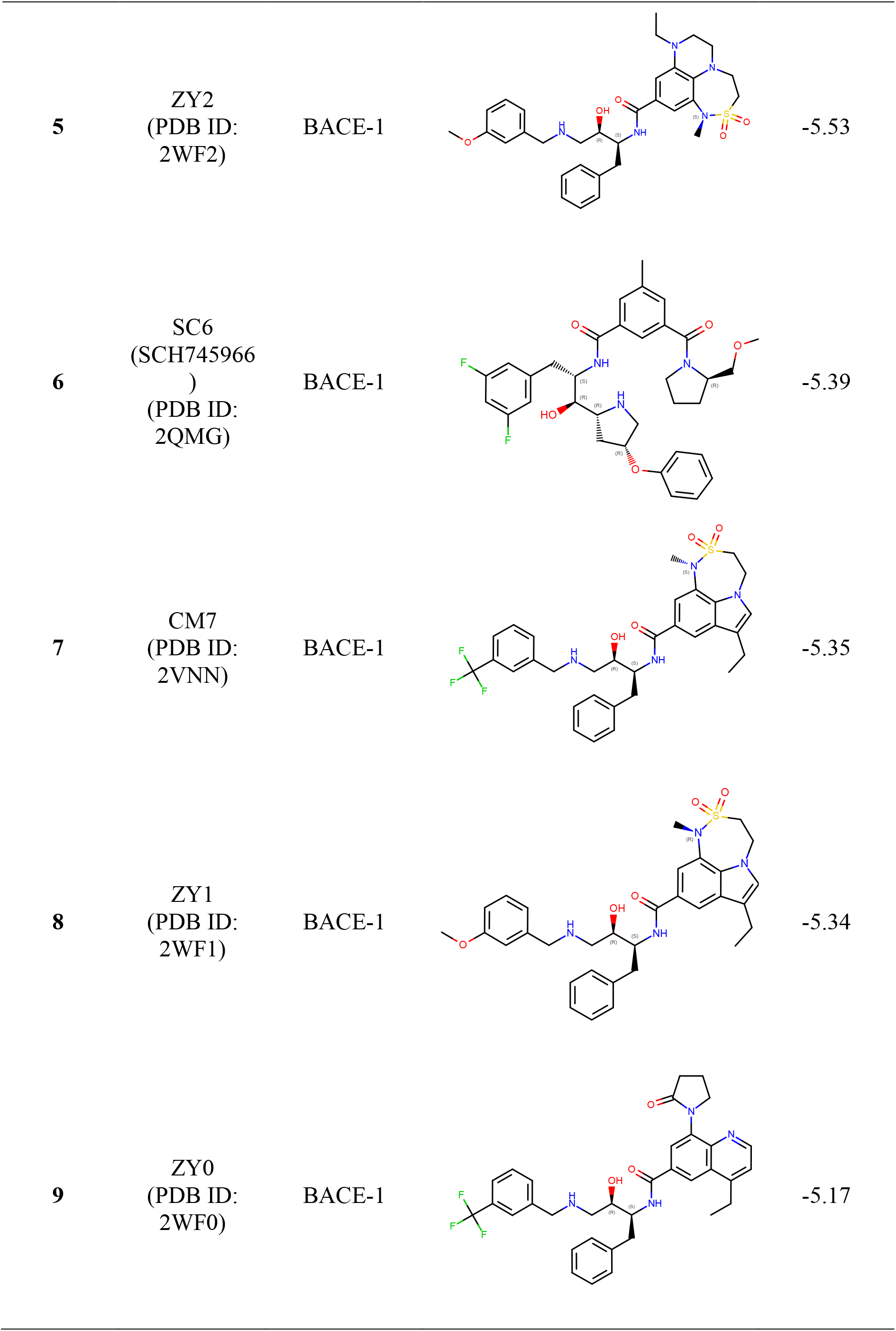
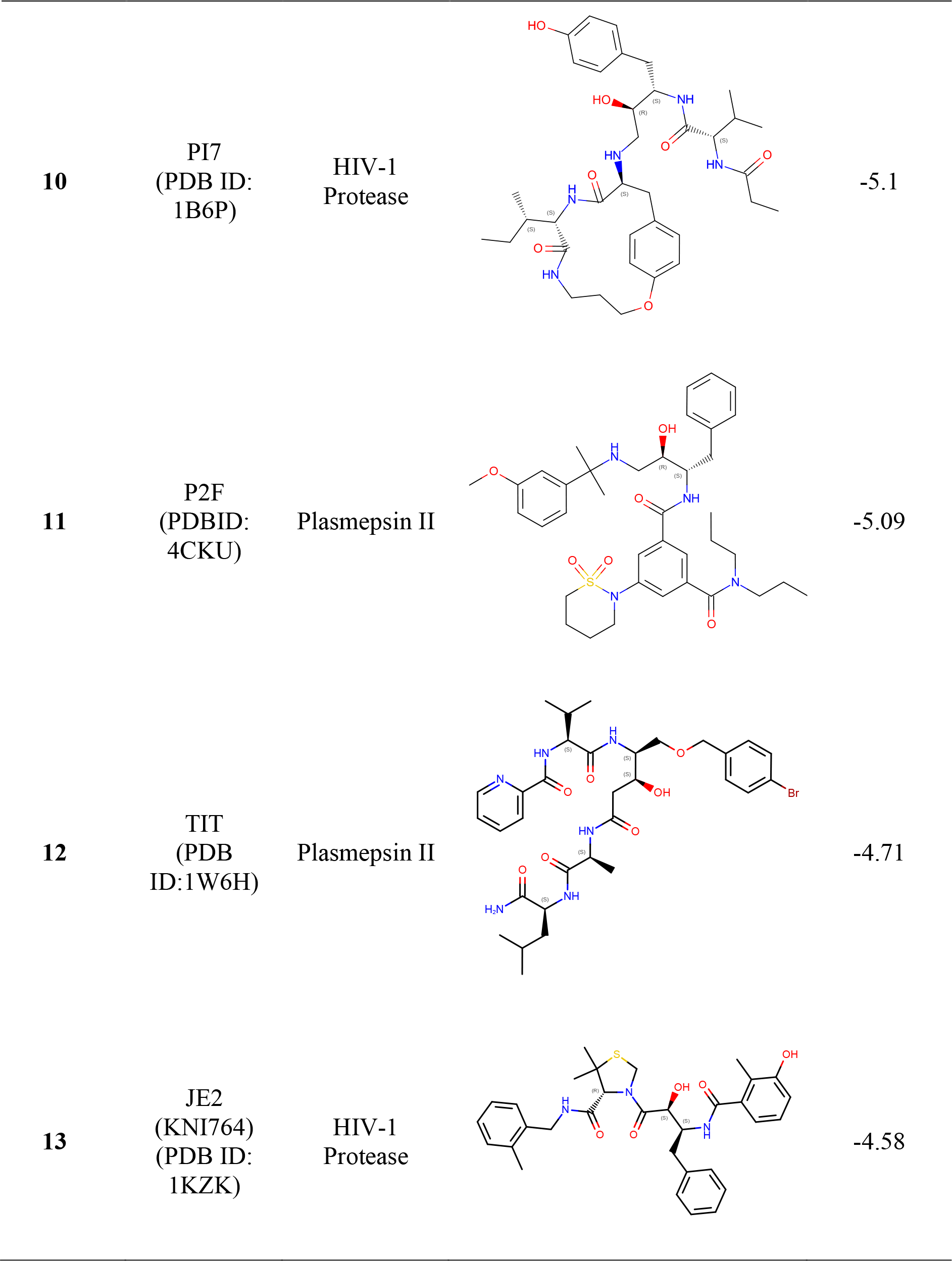
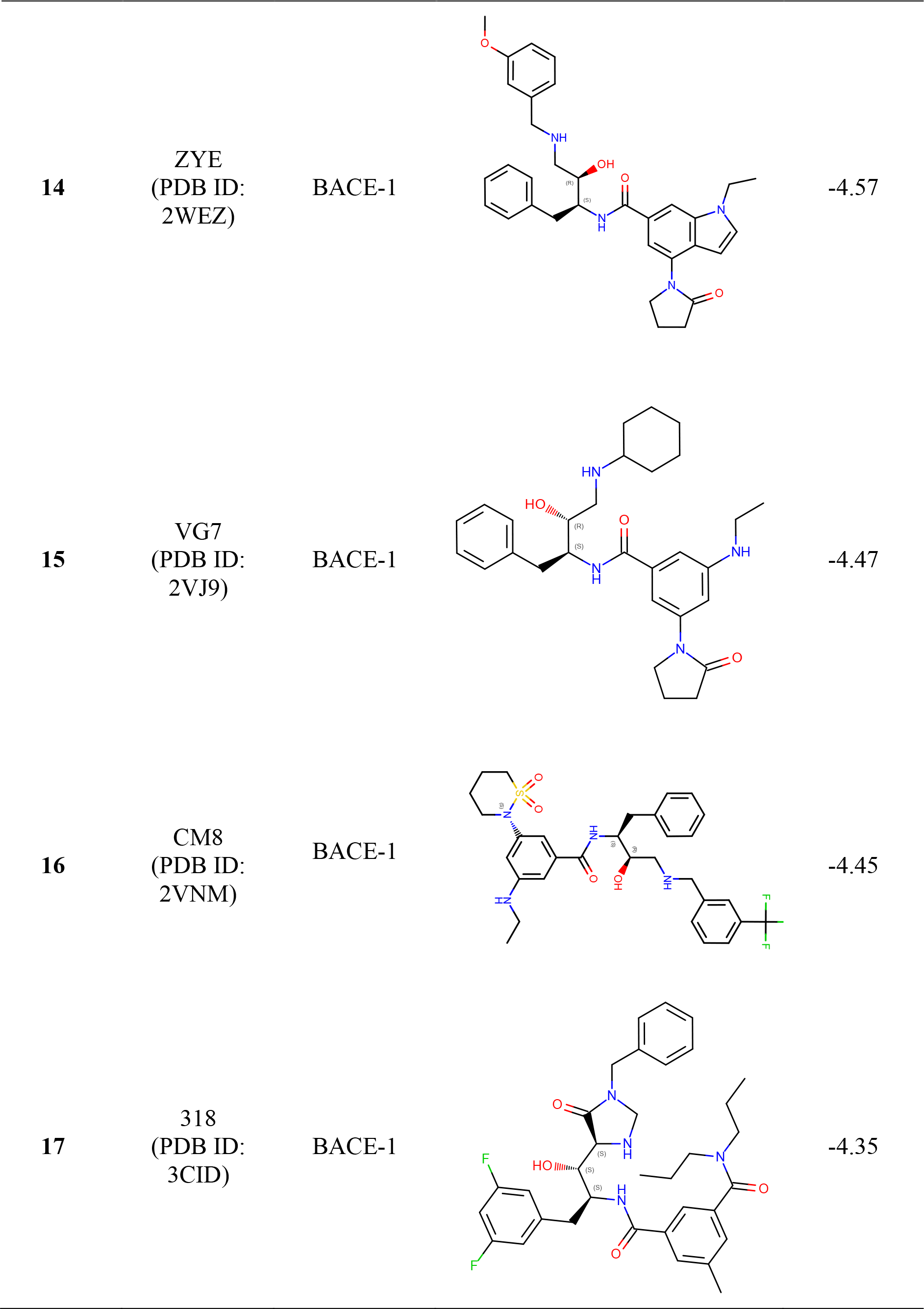
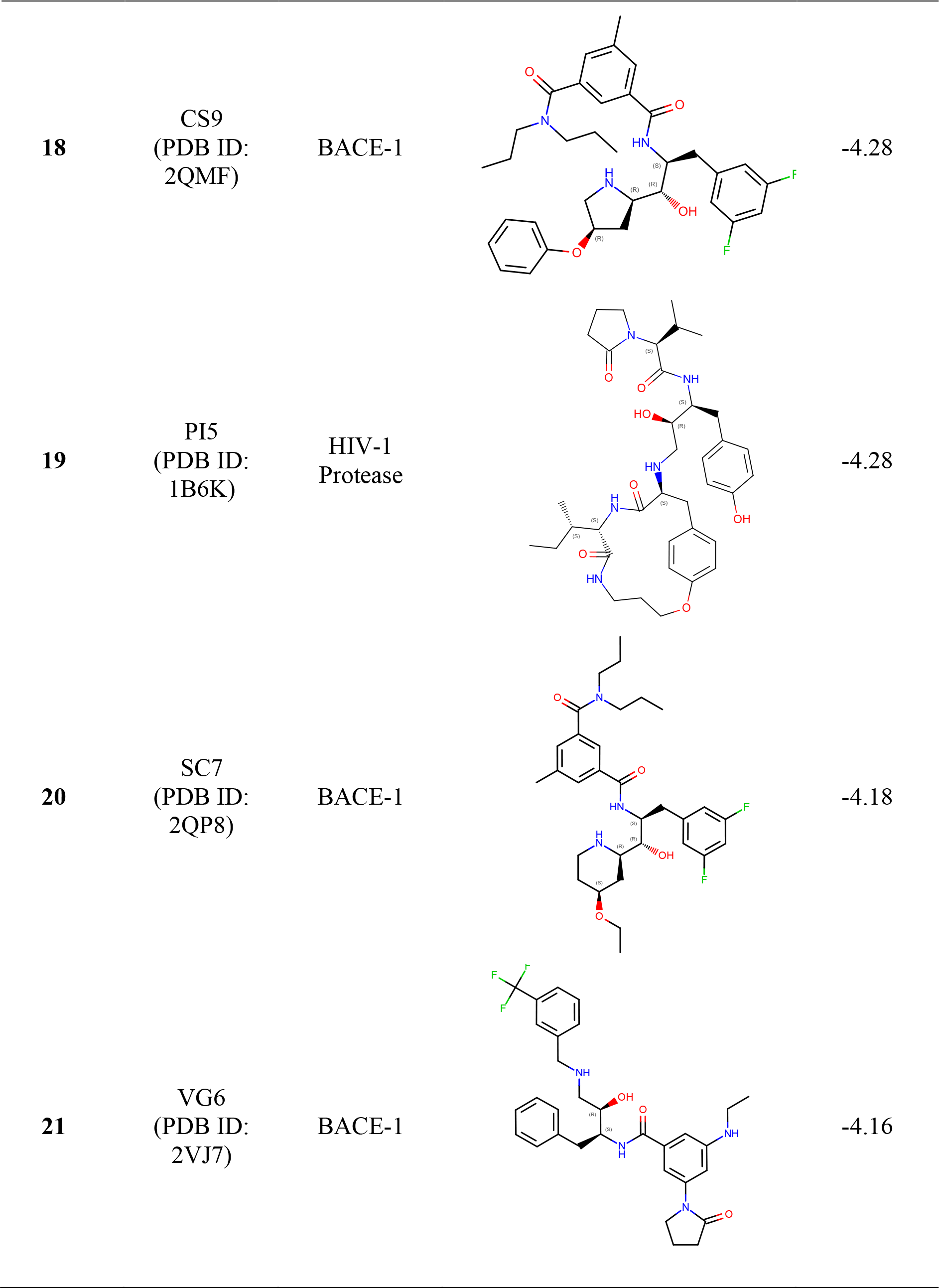
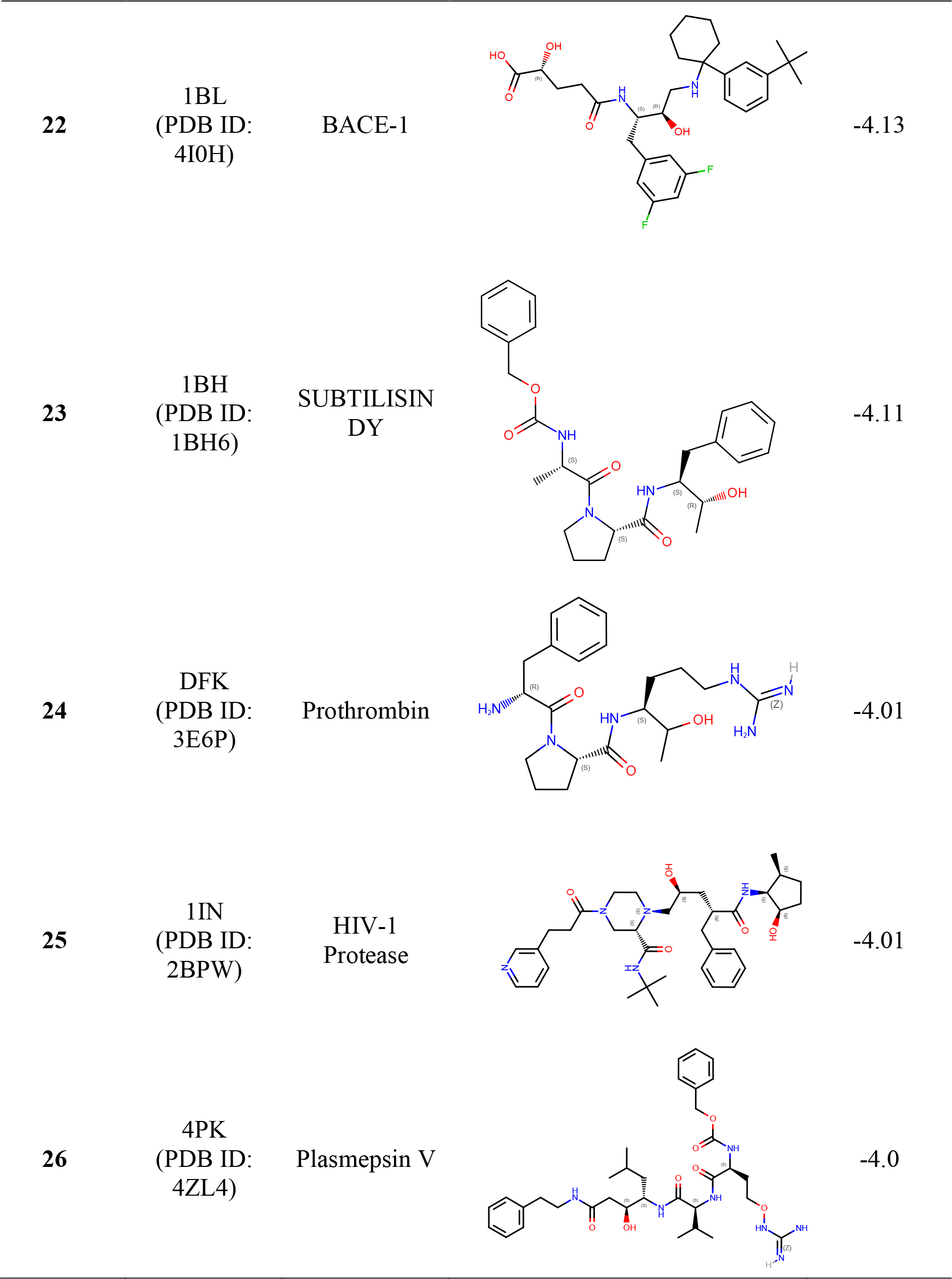
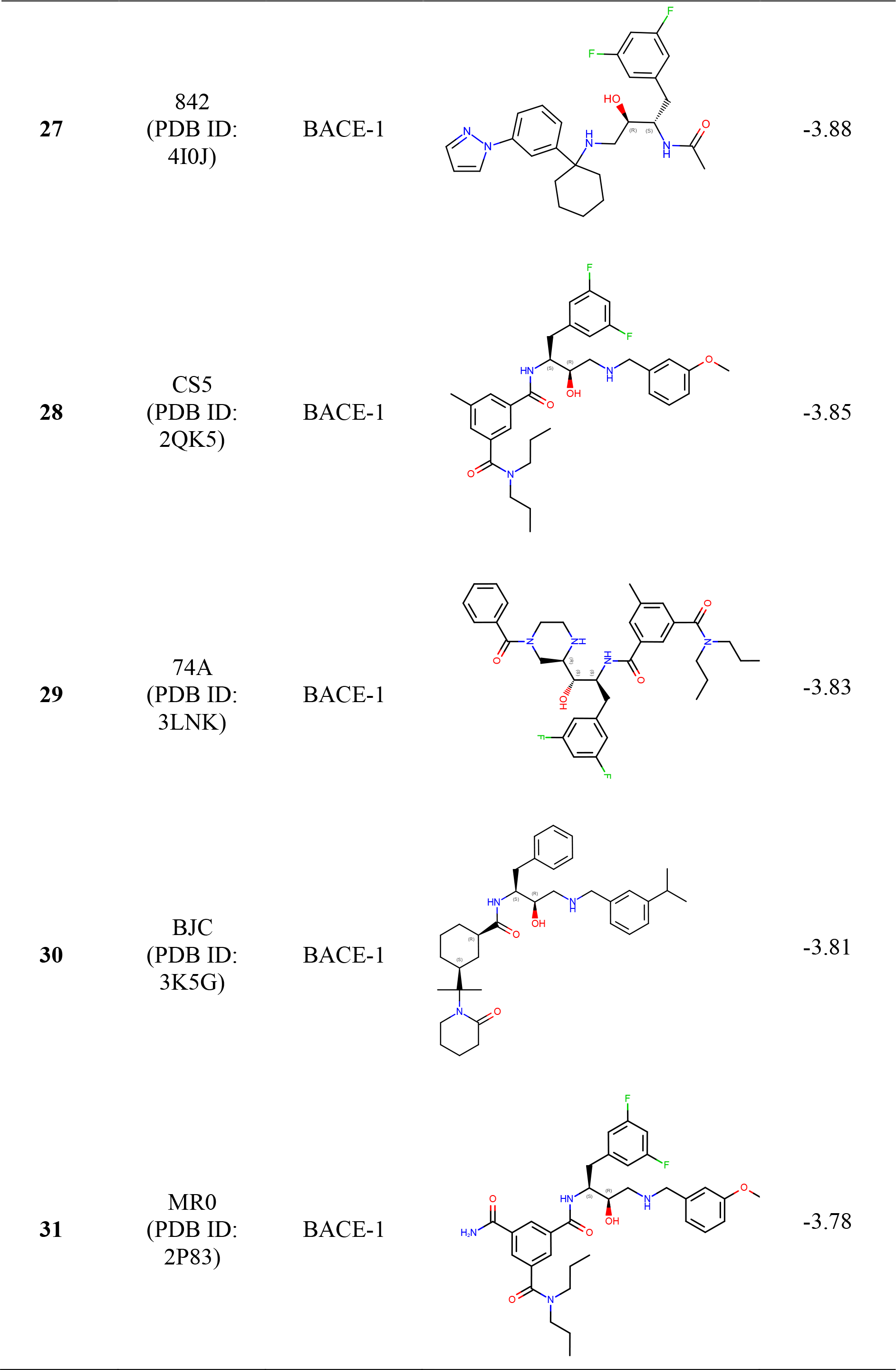
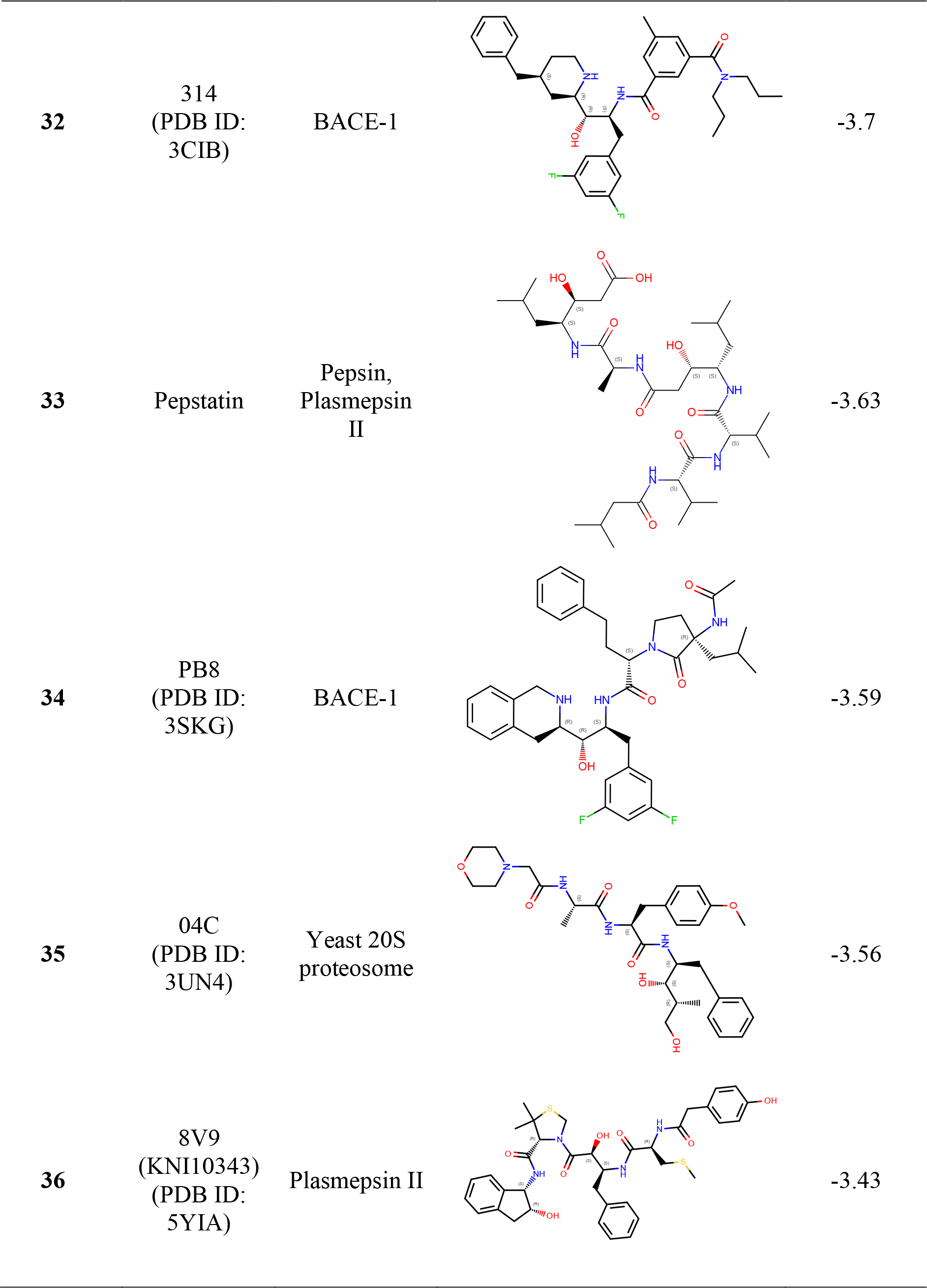
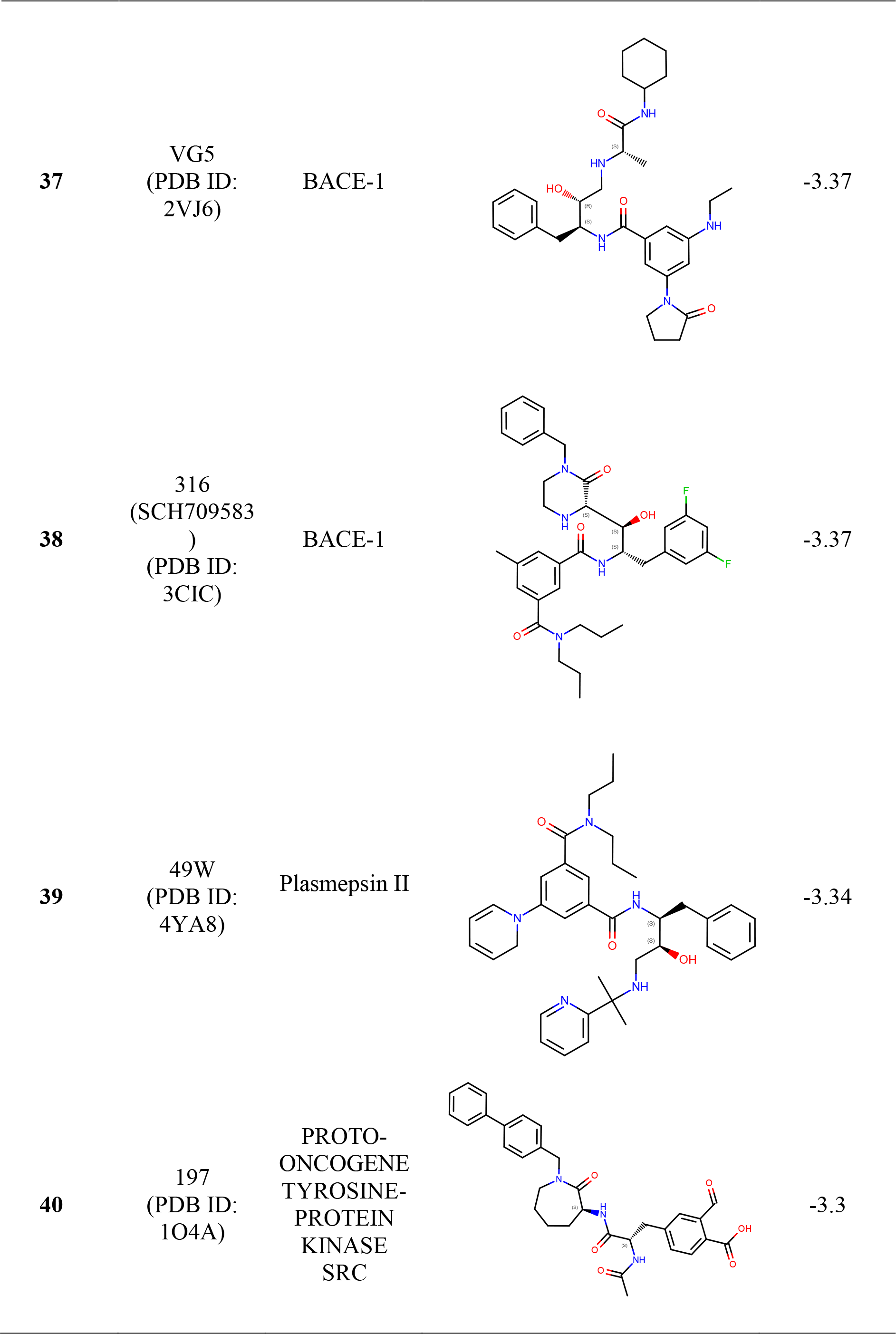
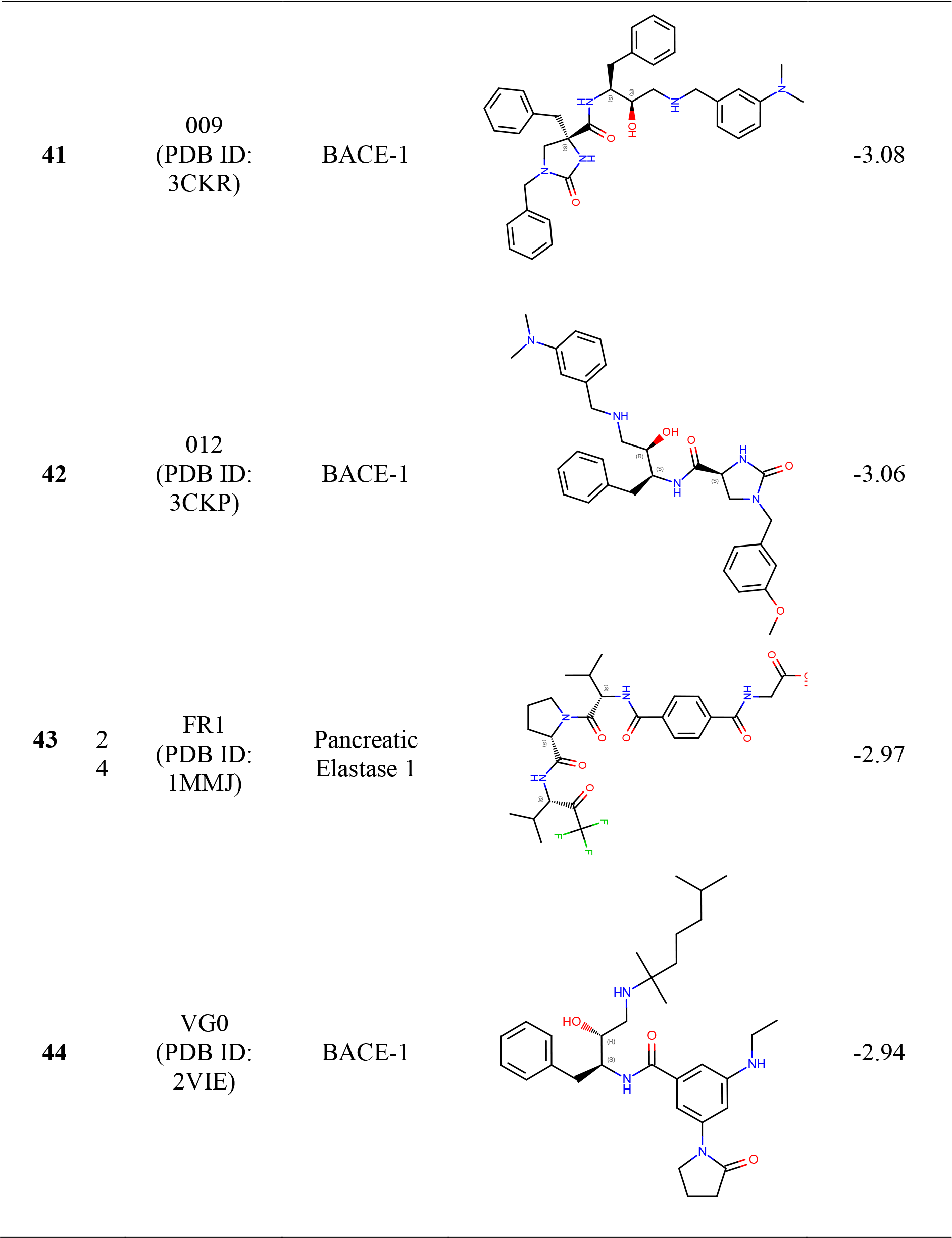
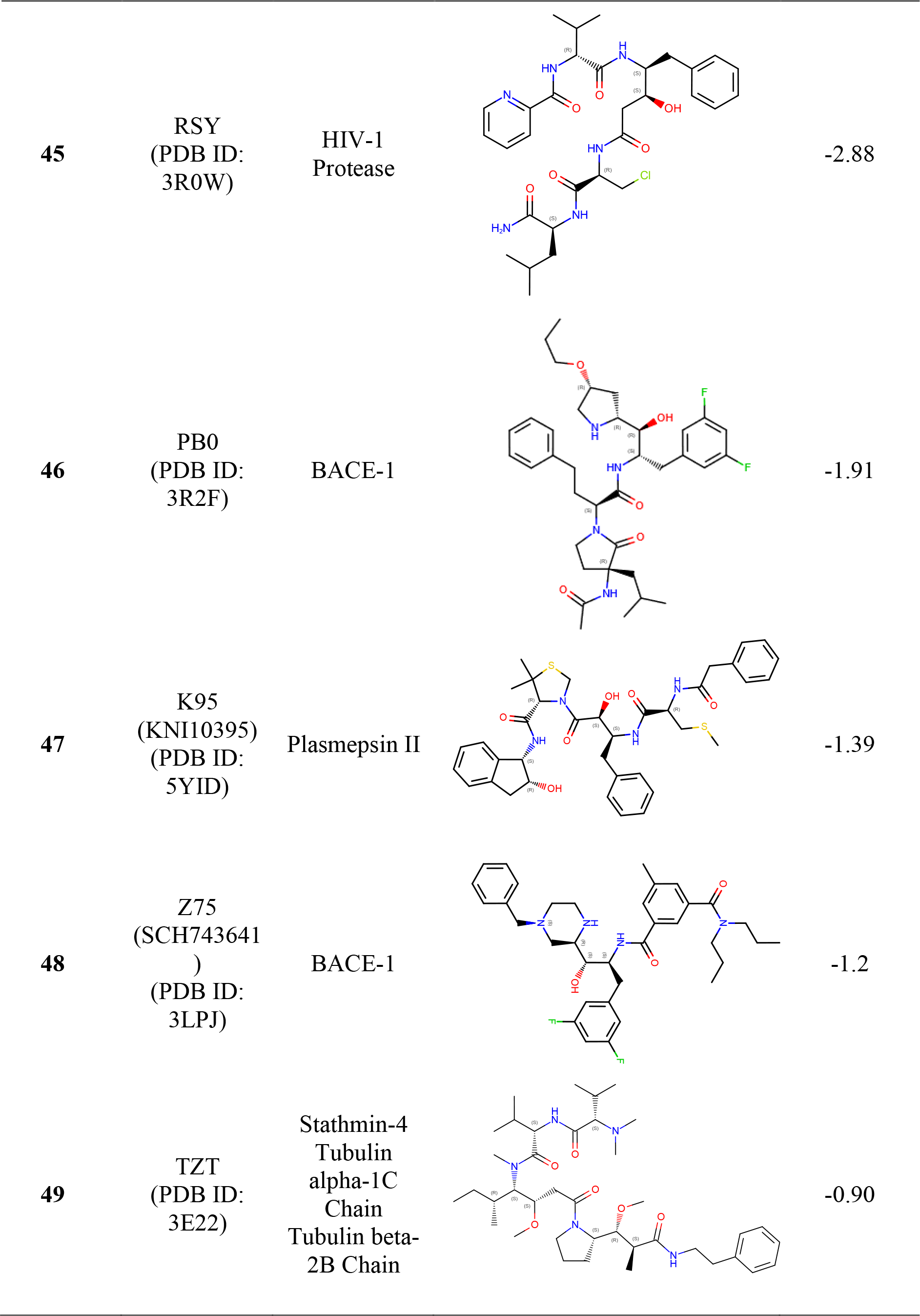
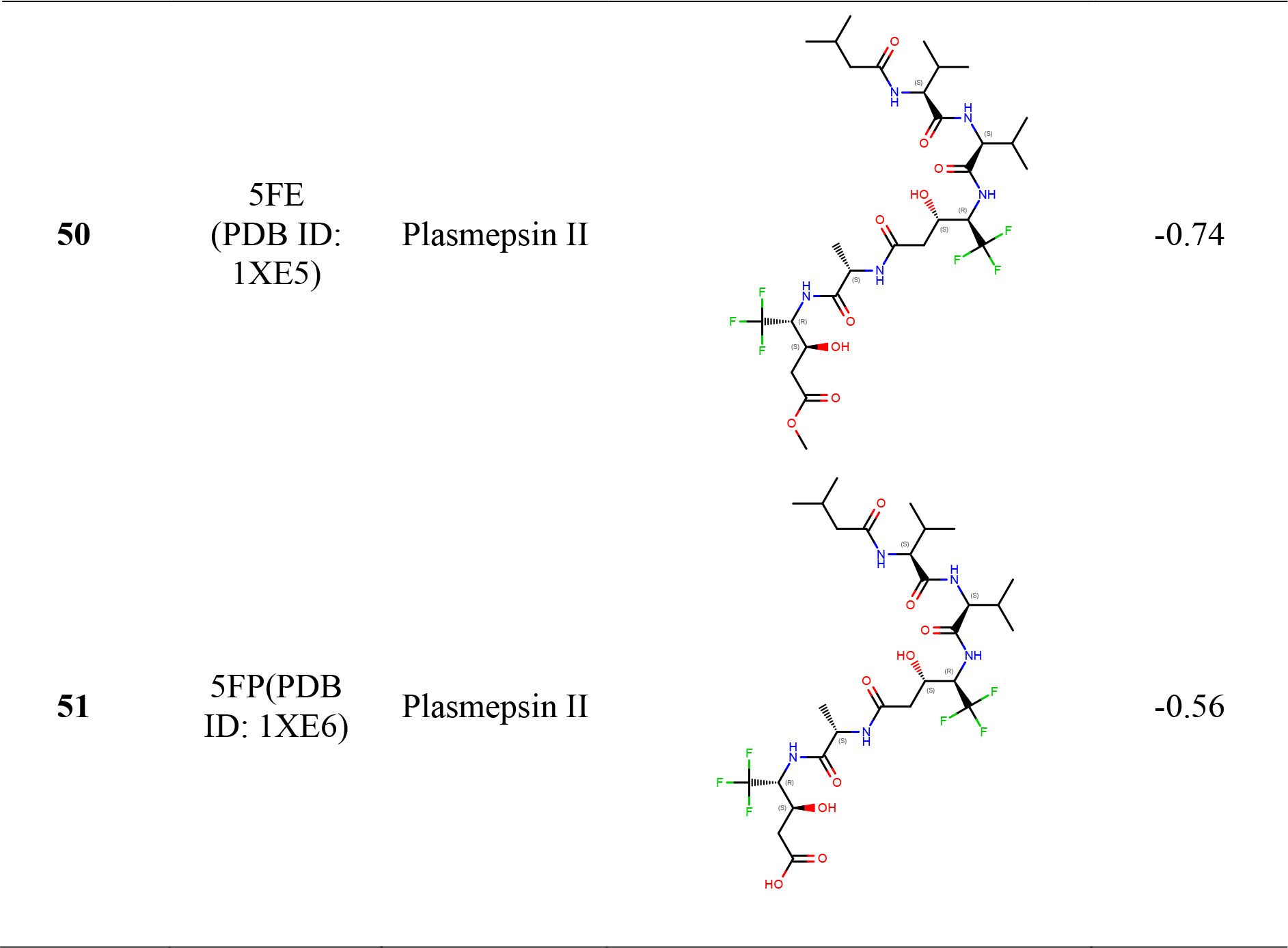
The inhibitors screened against *Tg*ASP5 active site. The ligand molecules are arranged based on docking score along with their covalent structures and PDB IDs from which they were extracted

